# Tree roots drive POM losses and transient MAOM gains across AM and ECM stands

**DOI:** 10.64898/2026.11.13.766288

**Authors:** Young E. Oh, Kurt A. Smemo, Sara M. Moledor, Andrew C. Eagar, Christopher B. Blackwood, Valentin Kurbel, Richard P. Phillips

## Abstract

Roots play a central role in soil organic matter (SOM) formation, stabilization and decomposition, yet how roots and soil properties modulate these dynamics is poorly understood. We used ingrowth cores to investigate how roots influence particulate organic matter (POM) and mineral-associated organic matter (MAOM) in forest plots dominated by arbuscular mycorrhizal (AM) and ectomycorrhizal (EcM) trees in Indiana (IN) and New York (NY), USA. Additionally, we amended a subset of cores with nutrients or water (controls) to examine how nutrient availability affects root-induced changes in POM and MAOM at a moderately N-limited site (IN) and a strongly N-limited site (NY). We tested three hypotheses: H1) root-induced changes in SOM fractions differ between tree-mycorrhizal types, with EcM roots promoting POM losses and AM roots enhancing MAOM gains, H2) increases in nutrient availability reduce POM loss and enhance MAOM formation, with the greatest changes occurring at the most N-limiting environments (NY), and H3) root-induced changes in POM and MAOM fluctuate over time, as initial gains in SOM are offset by SOM losses. We found partial support for H1 and H2 and modest support for H3. For H1, POM-C losses outpaced MAOM gains in EcM-dominated plots (as hypothesized) but both SOM fractions were mostly unchanged in AM-dominated plots. For H2, nitrogen addition did not increase POM-C (as hypothesized), but it did increase MAOM-C modestly in AM plots and EcM plots at the more N-limited (NY). In support of H3, MAOM gains observed after 24 months had generally diminished by 35 months. This was especially true for MAOM-N at the NY site which declined by 13.4%. Together, our results show that roots can simultaneously accelerate losses from POM and promote MAOM accumulation, and that tree-mycorrhizal type, environmental context and time modulate these effects. Additionally, our study suggests that short-term studies of root-SOM interactions may provide an incomplete view of soil C persistence drivers in forests, which can change dynamically over time.

## Introduction

The persistence of soil organic matter (SOM) determines how long carbon (C) remains in soils, yet the drivers of persistence remain debated (Scharlemann et al., 2014; Schmidt et al., 2011). Roots are often considered primary modulators of SOM dynamics (Rasse et al., 2005), influencing its formation (Sokol & Bradford, 2019), stabilization (Kravchenko et al., 2019) and turnover (Jastrow et al., 2007; Jilling et al., 2021). Living roots accelerate SOM decomposition by releasing labile rhizodeposits that stimulate extracellular enzyme production by microbes via ‘priming effects’ (Cheng et al., 2014; Finzi et al., 2015). Microbes can also slow SOM decomposition, as microbial necromass can sorb to reactive mineral surfaces (Liang et al., 2017; Sokol et al., 2019; Sokol & Bradford, 2019) and increase SOM persistence. Dead roots can affect SOM decomposition and persistence, as well. Likewise, decaying roots can increase microbial activity and turnover, leading to both priming effects and necromass stabilization on mineral surfaces (Beidler et al., 2023). Taken together, both living and dead roots can simultaneously *increase* and *decrease* SOM (Dijkstra et al., 2021), highlighting the need for carefully-designed field experiments where one or more factors can be controlled for (Dijkstra et al., 2021; Frey, 2019; Jones et al., 2009).

Tree species associating with different mycorrhizal partners possess a suite of traits (e.g., root inputs, mycorrhizal traits and litter quality; Keller et al., 2021; Lang et al., 2025; Phillips et al., 2013) that can influence root-SOM dynamics. Arbuscular mycorrhizal (AM) trees produce high-quality litters that lead to inorganic nutrient economies and diverse microbial communities whereas ectomycorrhizal (EcM) trees tend to produce slow cycling litters that promote organic nutrient economies and less diverse microbial assemblages (Eagar et al., 2022). How nutrient economies and soil microbial communities jointly regulate root-SOM dynamics is largely unknown (Hicks Pries et al., 2023; A. K. Lang et al., 2023). Actively growing EcM roots and EcM fungi can accelerate SOM decay by releasing exudates and extracellular enzymes to soil (Lv et al., 2026) or suppress litter decay (Beidler et al., 2021) by reducing the activity of saprotrophic fungi (Fernandez & Kennedy, 2016; Gadgil & Gadgil, 1971, 1975). AM roots also affect SOM pools by releasing exudates to soil (Lv et al., 2026) and by producing fast-decaying litters that can form putatively stable SOM (Beidler et al., 2023). Together, these findings suggest that both AM and EcM mycorrhizal roots can simultaneously build and deplete SOM, making their net effects difficult to predict without direct measurements of root-driven SOM gains and losses.

A second factor that can modulate the direction and magnitude of root effects on SOM is nutrient availability (Dijkstra et al., 2013). Forest soils amended with nitrogen (N) can increase SOM (Chen et al., 2018; Janssens et al., 2010) – changes that result, in part, from root and rhizosphere processes. In many soils, root-derived C stimulates microbes to make extracellular enzymes to release nutrients from SOM (Craine et al., 2007; Dijkstra et al., 2013) through microbial N mining (Moorhead and Sinsabaugh, 2006). Thus, if belowground C allocation is reduced in response to high soil N availability, SOM decay may be slowed (Liu & Greaver, 2010). Indeed, peroxidase-producing EcM fungi are often less abundant as inorganic N availability increases and more abundant as lignin-derived SOM increases (Argiroff et al., 2022; Clemmensen et al., 2015), suggesting that N-rich soils may experience less root-induced SOM decomposition (Lindahl et al., 2021). Alternatively, if high N availability limits belowground C allocation in plants, microbial growth and turnover may also decrease, reducing the potential for necromass stabilization. Resolving how changes in N availability affects SOM dynamics thus requires consideration of the direct and indirect effects of N availability on root and microbial activities.

The factors that control SOM dynamics and persistence also depend on the type of the SOM, as particulate (POM) and mineral-associated (MAOM) organic matter differ greatly in their turnover times and sensitivity to root-induced microbial activities (Lang et al., 2026). Most POM is considered unprotected and vulnerable to microbially-mediated decay whereas most MAOM is considered protected and insensitive to microbially-mediated decay (Kumar et al., 2020; Liu et al., 2026). Root exudates can alter both pools via changes in microbial activity (Chari and Taylor 2022). Exudates can decrease POM by providing microbes with an energy subsidy (i.e., priming effects; Kuzyakov 2010) or increase MAOM by facilitating the growth, turnover and stabilization of microbial necromass (Craig et al., 2022). Notably, some exudates (e.g., oxalic acid) can desorb and solubilize OM from minerals, leading to accelerated MAOM decay or to new MAOM formation as microbes die (Keiluweit et al., 2015). Thus, while root exudates are often believed to increase SOM losses owing to priming (Kuzyakov 2010), their net effects may be more variable owing to cascading effects on MAOM (Liu et al. 2026). Of course, POM and MAOM pools are not independent, as POM losses can lead to MAOM gains (Cotrufo et al., 2013; Dijkstra & Cheng, 2007; Lavallee et al., 2020; Ridgeway et al., 2022). Whether roots effects on POM and MAOM are convergent or divergent is largely unknown, necessitating studies of both pools under a range of environmental conditions (Jackson et al., 2017; Keller et al., 2021; A. Lang et al., 2026).

The positive relationship between root density and SOM stocks is often cited as evidence that roots control SOM dynamics (Heckman et al., 2023; Malhotra et al., 2025). However, static relationships likely yield an incomplete picture about the processes that control soil C persistence. In this study, we used a multi-year root-ingrowth core experiment across U.S. eastern temperate forests to determine how roots influence SOM dynamics and whether these effects depend on tree-mycorrhizal association, environmental context and nutrient availability. We addressed three questions. First, how does root ingrowth alter SOM C (and N) pools, and do these responses vary between AM- and EcM-dominated forests and across sites? We hypothesized that root-derived SOM-C changes would differ between tree mycorrhizal dominance as EcM roots promote POM loss and AM roots enhance MAOM gains. We further hypothesized that these responses would vary among sites because environmental and edaphic conditions modify microbial constraints on SOM decomposition and formation. Second, to what extent does nutrient availability alter relationships between roots and SOM, and do these responses depend on mycorrhizal type and site context? We hypothesized that alleviating nutrient limitation (by fertilizing ingrowth cores), would alter root–SOM relationships by changing nutrient-acquisition and SOM-processing, with stronger responses occurring in the most N-limited plots (e.g., EcM-dominated forests in NY). Finally, we asked whether changes in POM and MAOM observed in the short-term (two years) persisted, accumulated or diminished with time (after three years). Together, these approaches allowed us to evaluate how tree-mycorrhizal association, nutrient availability, environmental context, and time interact to determine the consequences of root colonization for SOM persistence.

## Methods

### Experimental design

We conducted a root ingrowth-core experiment to examine how root activity and nutrient availability shape SOM dynamics across forests differing in climate, tree-mycorrhizal dominance and edaphic characteristics. Ingrowth cores were installed in 2018 or 2019 in arbuscular mycorrhizal (AM)- and ectomycorrhizal (EcM)-dominated plots in secondary growth forests in Indiana (IN) and New York (NY), USA. At each site, plots were selected along a gradient in tree-mycorrhizal dominance - arbuscular mycorrhizal (AM)- or ectomycorrhizal (EcM)-dominated stands - based on the relative basal area of canopy trees associated with each mycorrhizal type (across both sites, AM trees were 83% of the basal area in AM plots, EcM trees were 91% in EcM plots). At IN, five 20 × 20 m plots were established for each tree mycorrhizal type (Table 1). At NY, nine 15-m-radius plots of each mycorrhizal type were established, with three replicate plots across three regions (Table 1). Within each plot, cores were assigned to either a fertilization treatment, receiving a dissolved nutrient mixture bimonthly, or a control treatment receiving an equivalent volume of deionized water. Cores were collected after 24 months and 35 months to quantify root biomass and morphology, SOM C and N pools, potential extracellular enzyme activities, and soil N cycling. Additional cores were harvested after 35 months to quantify root biomass and morphology, and SOM C and N pools.

**Table 1).**
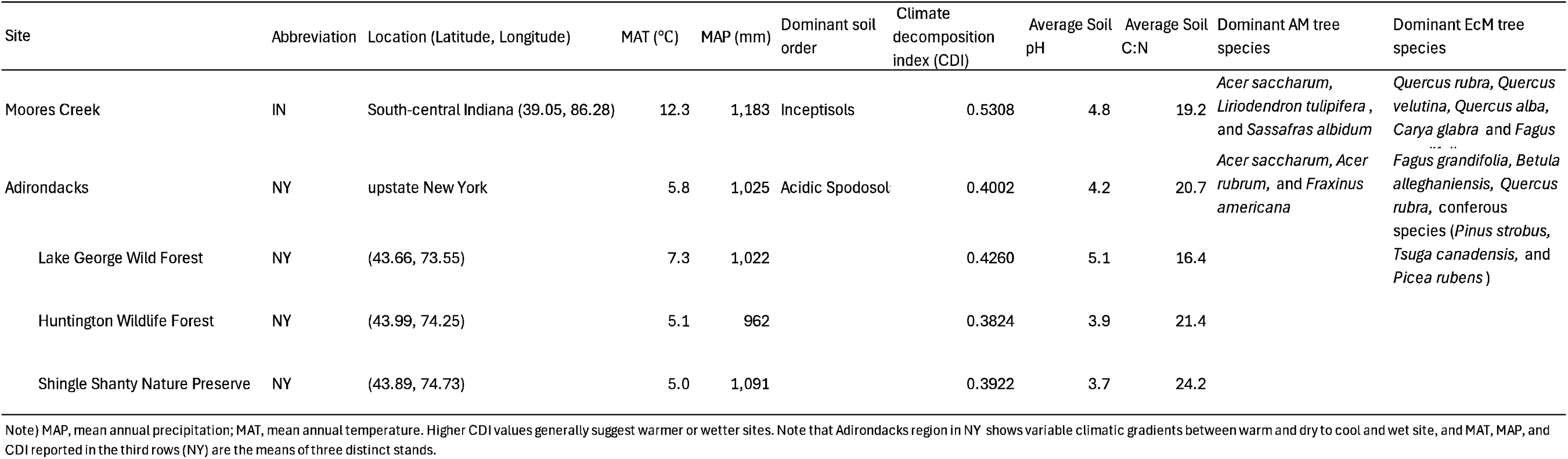
Location of study sites with mean climate conditions, soil order and dominant AM and ECM tree species. Climate decomposition index (CDI) was the means of the monthly-scale CDIs is during 2019 - 2021. The CDIs is were calculated using the climatic data from PRISM Climate Group(PRISM Climate Group 2021), following the formula suggested by Adair et al. (2008) and Zhou et al. (2015).

| Site | Abbreviation | Location (Latitude, Longitude) | MAT (°C) | MAP (mm) | Dominant soil order | Climate decomposition index (CDI) | Average Soil pH | Average Soil C:N | Dominant AM tree species | Dominant EcM tree species |
|---|---|---|---|---|---|---|---|---|---|---|
| Moores Creek | IN | South-central Indiana (39.05, 86.28) | 12.3 | 1,183 | Inceptisols | 0.5308 | 4.8 | 19.2 | *Acer saccharum*, *Liriodendron tulipifera* , and *Sassafras albidum* | *Quercus rubra*, *Quercus velutina*, *Quercus alba*, *Carya glabra* and *Fagus* [illegible] |
| Adirondacks | NY | upstate New York | 5.8 | 1,025 | Acidic Spodosol[illegible] | 0.4002 | 4.2 | 20.7 | *Acer saccharum*, *Acer rubrum*, and *Fraxinus americana* | *Fagus grandifolia*, *Betula alleghaniensis*, *Quercus rubra*, conferous species (*Pinus strobus*, *Tsuga canadensis*, and *Picea rubens* ) |
| Lake George Wild Forest | NY | (43.66, 73.55) | 7.3 | 1,022 |  | 0.4260 | 5.1 | 16.4 |  |  |
| Huntington Wildlife Forest | NY | (43.99, 74.25) | 5.1 | 962 |  | 0.3824 | 3.9 | 21.4 |  |  |
| Shingle Shanty Nature Preserve | NY | (43.89, 74.73) | 5.0 | 1,091 |  | 0.3922 | 3.7 | 24.2 |  |  |

Note) MAP, mean annual precipitation; MAT, mean annual temperature. Higher CDI values generally suggest warmer or wetter sites. Note that Adirondacks region in NY shows variable climatic gradients between warm and dry to cool and wet site, and MAT, MAP, and CDI reported in the third rows (NY) are the means of three distinct stands.

### Study sites

Moores Creek is an 85 yr-old hardwood forest in south central Indiana (39.983° N, 90.817° W) and is part of the IU Research and Teaching Preserve (Table 1). Overstory composition is typical of temperate broadleaf forests and includes *Quercus spp., Carya spp., Acer saccharum, Fagus grandifolia* and *Liriodendron tulipifera*. For the timeframe of this study (2019–2021), mean annual temperature was 12.3 °C mean annual precipitation was 1,183 mm yr ¹, respectively, based on PRISM climate data (PRISM Climate Group, 2021). Soils at IN are predominantly Inceptisols. More details and the tree communities and soils are available in Rosling et al. (2016), Midgley et al. (2019) and Mushinski et al. (2019).

The three forests in the Adirondack (ADK) region of New York (NY) include Lake George Wild Forest (DL; 43.661° N, 73.545° W), Shingle Shanty Preserve (SS; 43.894° N, 74.732° W) and Huntington Wildlife Forest (HF; 43.987° N, 74.245° W). Tree species composition at NY included several species also occurring at IN, but EcM-dominated stands additionally contained conifers such as *Pinus strobus, Tsuga canadensis* and *Picea rubens* (Table 1). Mean annual temperature was 5.8 °C and mean annual precipitation was 1,025 mm yr ¹ for the time period of this study. Soils at the NY sites are predominantly acidic Spodosols and detailed information on these sites can be found in Eagar et al (2023) and Cortese et al (2025).

### Ingrowth core and experimental treatments

At each plot, we inserted two cylindrical root ingrowth cores (5.77-cm inner diameter × 15-cm height) into soil. The cores were constructed from rigid plastic mesh with 3-mm openings that allows roots and associated mycorrhizal fungi to colonize the cores. Before core installation, the Oi horizon was removed, and soil was collected from each core location to characterize initial soil conditions (“initial soil” hereafter). Soils collected adjacent to the core locations were homogenized across plots within each mycorrhizal type and used to fill the ingrowth cores.

At IN, homogenized soil was mixed with autoclaved sand at a 2:1 soil ratio by dry mass to improve drainage, while no sand was added to the soil mix at NY. The sand used at IN contained CaCO and consequently the pH of the sand-soil mix was elevated by ∼2 pH units relative to the surrounding mineral soil (Fig. S16; S17; Table 1). Because pH can alter microbial activity and SOM dynamics (Heckman et al., 2023), we compared our results with those from an independent ingrowth-core experiment conducted in the same plots using non-carbonate sand, which did not alter soil pH. Despite differences in soil pH between the experiments, we found broadly comparable patterns of root ingrowth and changes in SOM pools over time (Fig. S17), suggesting that the patterns observed in our study were driven more by root-microbe interactions than by soil pH.

A subset of ingrowth cores were assigned to one of two nutrient treatments: fertilization with a dissolved nutrient mixture or a control receiving an equivalent volume of deionized water. Treatments were applied to cores (via injection) bimonthly during the growing season (May to October) for two consecutive years. Fertilization treatment (50mL per core) included 198g of Ca(NO_3_)2-(H_2_O)4 (solution), 103g of KH_2_PO_4_ (solution), 156g of Osmocote Plus 15-9-12 (including S, Mg, Cu, Fe, Mn, Mo, and Zn). Control cores received the same volume of deionized water without added nutrients. Likewise a subset of unfertilized ingrowth cores were left in the ground for 35 months to explore changes in root-induced POM and MAOM pools over time.

### Soil organic matter analyses

The ingrowth cores were collected in 2021, and soils and roots were separated after passing soils through a 2mm sieve. Roots were oven-dried at 60 for two days and soils were air-dried (> one week). Air-dried soils were separated into three fractions based on density (Sollins et al., 2009) using sodium polytungstate (SPT), which has a density of 1.71 g/mL. To isolate free particulate organic matter (fPOM, hereafter), we subsampled 3g of soil and submerged in SPT solution and centrifuged at 3700rpm for 30 minutes. The remaining soil was mixed with plastic beads and shaken for 16 h at 200 oscillations per minute to disrupt soil aggregates. The samples were then centrifuged to isolate organic matter less than the density of SPT, which consists of occluded particulate organic matter (oPOM, hereafter). The remaining organic matter was considered mineral-associated organic matter (MOAM). All fractions of organic matter were rinsed with Nanopure water to remove residual SPT. Samples with recovery rates ranging 99-102% (indicating residual SPT) were re-rinsed. All soil fractions were oven dried at 65 and analyzed for percent C and N using an elemental combustion system (Costech Analytical Technologies).

To express C and N contents of each soil organic matter (SOM) fraction (fPOM, oPOM, and MAOM) on a common soil-mass basis, we calculated fraction C and N concentration (mg C or N g ¹ soil) by multiplying the dry mass of each recovered fraction by its measured C or N concentration and dividing by the dry mass of soil used for fractionation. Given that soil used for the IN cores was mixed with sand at a 2:1 soil:sand ratio (by dry mass), we corrected the C & N percent concentration from the elemental combustion system and grams of soil samples used for fractionation in calculating C and N concentration in MAOM fraction.

### Root trait measurements

For each core, any coarse and dead roots were separated out, fine roots were oven-dried at 60° C for a minimum of 24 hours, then compiled and weighed for total (fine) root biomass per core. The total root biomass data was standardized by dividing the biomass by the total dry soil weight per core (‘root density’, hereafter) to account for root biomass differences in cores that lost soil via erosion, etc. For root trait data, up to five root segments representative of the overall core demographic were selected per core and scanned and analyzed using WinRhizo software (Régent Instruments Inc., Canada). First three root segments were considered ‘absorptive root’, and absorptive root density (mg of roots g^-1^ soil) was calculated based on the root density. Measured root traits included: 1) specific root length (SRL): total root length (cm) x mass^-1^ (g), 2) specific root surface area (SRA): surface area (mm^2^) x mass^-1^ (g), 3) root tissue density (RTD): weight (g) x volume^-1^ (mm³), 4) specific root tip abundance (SRTA): # of root tips x mass^-1^ (g), 5) branching intensity (BI): # of root tips x length^-1^ (cm) and 6) mean root diameter (cm). To avoid faulty estimates in WinRhizo’s calculations (discussed in Rose (2017)), SRL, SRA, and RTD were calculated using respective lengths, surface areas, and volumes summed across all diameter classes.

### Statistical analyses

We used a mixed-effect model (‘lme4::lmer’ via REML) to evaluate the effects of root ingrowth, nutrient addition, tree-mycorrhizal dominance and site on changes in C and N concentration each soil organic matter fraction (e.g., fPOM, oPOM, and MAOM), with plot included as a random effect when multiple observations were associated with the same plot. For analyses in which observations were independent at the plot level and no additional grouping structure required a random effect, ordinary linear models were fitted using the ‘lm’ function.

To assess root-induced changes on SOM C and N concentrations, fPOM, oPOM and MAOM C and N concentrations from the initial soil concentration was subtracted from the fPOM, oPOM and MAOM C and N concentration measured at the end of the ingrowth period. We refer to these changes over time as ΔfPOM-C&N, ΔoPOM-C&N and ΔMAOM-C&N, where positive values indicate net accumulation in each fraction relative to initial soil and negative values indicate net loss. We compared ΔfPOM-C&N, ΔoPOM-C&N and ΔMAOM-C&N between AM- and EcM-dominated plots at both sites using ‘lme4::lmer’ via REML. Site, tree-mycorrhizal dominance and their interactions were included as fixed effects. We also tested whether the relationship between absorptive root density (mg biomass * g^-1^ soil) and ΔSOM differs among sites or between AM-vs EcM dominated plots.

To test whether nutrient addition altered POM- and MAOM-C and N responses and whether treatment effects differed between sites or mycorrhizal types, fertilized and control cores were compared within plots, using mixed-effects models. We additionally tested whether nutrient addition modified relationships between absorptive root density and changes in C and N in the SOM fractions (ΔSOM) by including the interaction between nutrient treatment and absorptive root density. When appropriate, relationships were evaluated separately among treatment, site, or tree-mycorrhizal groups.

To compare the responses among variables to nutrient addition across different measurement units, we additionally calculated standardized fertilization effect sizes separately for each site and mycorrhizal type (Fig. S19, S20). The mean paired difference was then standardized by the standard deviation of the paired differences across plots (mean ΔFert / S.D. of ΔFert), where positive mean differences indicate greater values under fertilization, whereas negative effect sizes indicate lower values under fertilization relative to controls. The analyses were applied to potential extracellular enzyme activities (acid phosphatase, N-acetyl-β-glucosaminidase, β-glucosidase, phenol oxidase, and peroxidase), N mineralization and net nitrification rates, and root-trait and changes in POM-C, MAOM-C, and MAOM-C:N. For each standardized effect size, error bars were shown using ‘approximate 95% confidence intervals’ calculated as the standardized mean paired difference ± 1.96 times its standardized standard error (a simple normal-approximation interval approach). Significance of fertilization effects was evaluated using two-sided paired t-tests comparing fertilized and control cores within plots. Tests were conducted separately for each site and mycorrhizal type.

To examine temporal changes in SOM following continued root ingrowth, POM- and MAOM-C and N concentrations measured after 24 and 35 months were compared with initial-soil values and between sampling periods. The mixed-effect model included experimental duration, site, and mycorrhizal dominance as fixed effects, with plot included as a random effect to account for repeated observations associated with the same plots.

To assess how root trait organization varied between initial soils and roots collected after two years of ingrowth, we performed principal component analysis (PCA). PCA was conducted using seven fine-root traits: root diameter, specific root length (SRL), specific root area (SRA), specific root tip area (SRTA), root tissue density (RTD), branching intensity (BI), and root density. Root traits were standardized prior to PCA to account for differences in measurement units and variation among traits. We first performed PCA across all observations to characterize overall root trait organization and evaluated differences between mycorrhizal types (AM vs. EcM) and sampling times (initial soils vs. 24-month ingrowth cores) using permutational multivariate analysis of variance (PERMANOVA; vegan::adonis2), with site included as a covariate (Fig. S5) and assessed differences in data dispersion using homogeneity of multivariate dispersion using vegan::betadisper (Fig. S5). We then generated separate PCAs for AM- and EcM-dominated plots to examine mycorrhizal-specific patterns of root trait coordination and their relationships with SOM-C dynamics (Fig. 3). For each mycorrhizal type, we examined associations between the dominant principal components (PC1 & PC2) and SOM-C pools in initial soils or changes in particulate organic matter C (ΔPOM-C) and mineral-associated organic matter C (ΔMAOM-C) after two years of root ingrowth (Fig. 3). Changes in SOM-C were calculated relative to the corresponding initial soil values. To identify individual root traits associated with variation in SOM-C, we additionally performed redundancy analysis (RDA; vegan::rda) with stepwise model selection (vegan::ordistep). Relationships between selected root traits and changes in SOM-C were subsequently evaluated using linear regression models.

To report the significance of each nested group in the model after accounting for all other groups, Type III ANOVA with Satterthwaite’s Method using the ‘lme4::anova’ was performed to summarize the results of each model and report in the corresponding figures. When necessary, estimated means and pairwise contrasts were calculated using the ‘emmeans::emmeans’, with Tukey adjustment for multiple comparisons. Model assumptions were evaluated by inspecting residual and quantile–quantile plots. Statistical significance was assessed at α = 0.05. All statistical analyses were conducted in R (R Core Team, 2025).

## Results

### The effects of root ingrowth on SOM fractions after 24 months

While root density was similar between AM vs EcM plots (Fig. 2d), changes in the distribution of C among SOM fractions relative to initial soils varied between tree mycorrhizal types. In EcM plots, free light POM-C (fPOM-C) and occluded POM-C (oPOM-C) declined by 69.1% and 47%, respectively, after two years (Fig 1a, 1b; Table 2, S6). In AM plots, fPOM-C and oPOM-C declined by approximately 46% and 23%, respectively (Table 2, S6). The decline in oPOM-C was significantly greater in EcM than AM plots at IN (β = −7.32, P = 0.032), whereas the corresponding difference in fPOM-C was not statistically significant (P = 0.219). Changes in POM-C were generally larger than those in MAOM-C, contributing to similar patterns in total SOM-C (Fig. S1). Likewise, we observed similar patterns in the distribution of N among the three SOM fractions (Fig. S3).

**Fig. 1).**
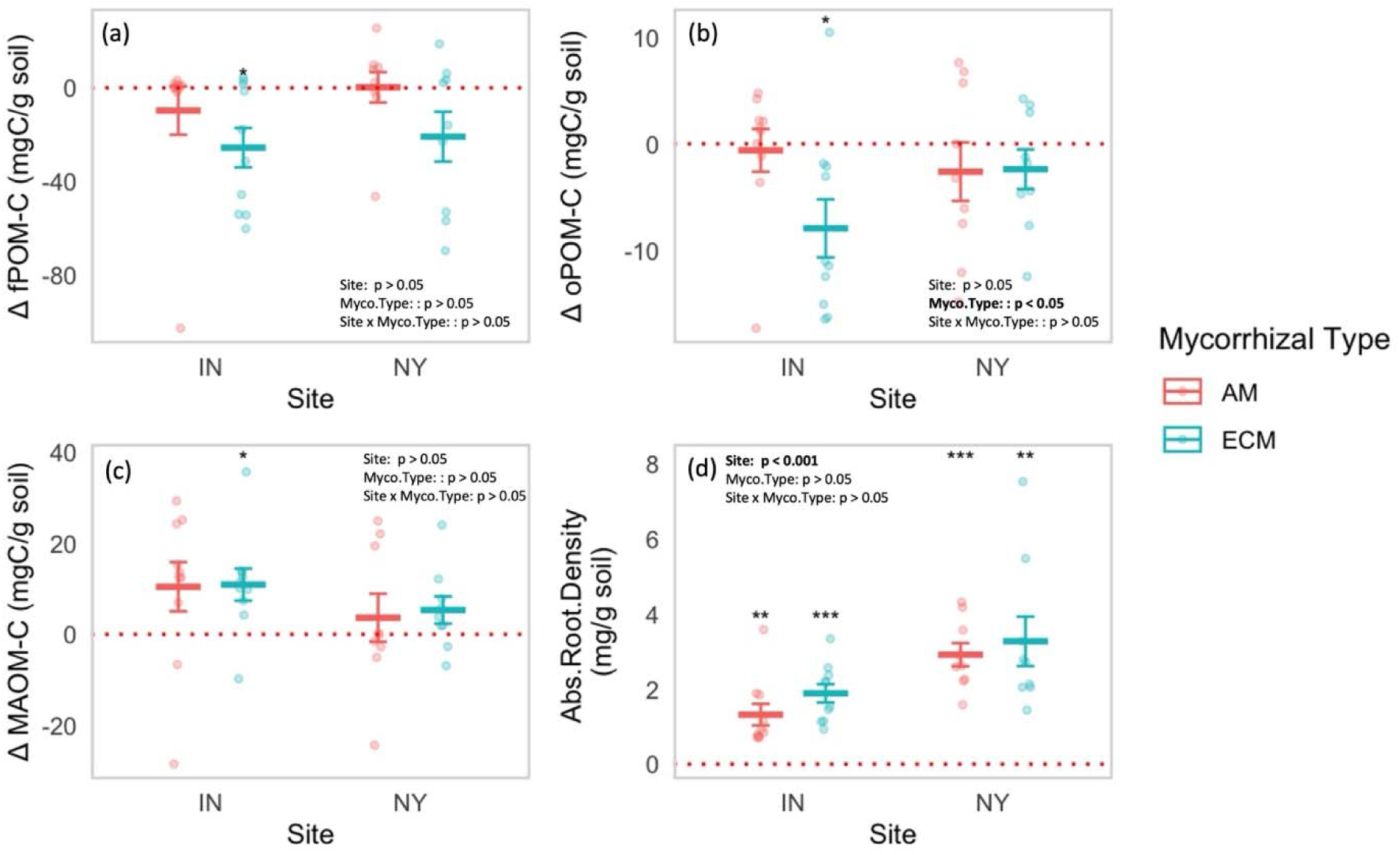
The effects of Site, mycorrhizal type, and their interaction on changes in soil organic matter C concentrations relative to initial soils (ΔSOM-C; mg C g ¹ soil) in the (a) free particulate organic matter (fPOM), (b) occluded particulate organic matter (oPOM), (c) mineral-associated organic matter (MAOM) fractions, and (d) absorptive root density (mg root g ¹ soil) in 24 months of new root ingrowth. Points represent individual observations, and crossbars indicate the mean ± SE for each Site × mycorrhizal type combination. Red dotted horizontal lines denote no change relative to initial soil (Δ = 0). Fixed effects of Site, mycorrhizal type, and their interaction were evaluated using linear models (lm). Asterisks indicate whether ΔSOM-C differed significantly from zero within each Site × mycorrhizal type combination (P < 0.05, P < 0.01), while model summaries are shown within each panel.

**Fig. 2).**
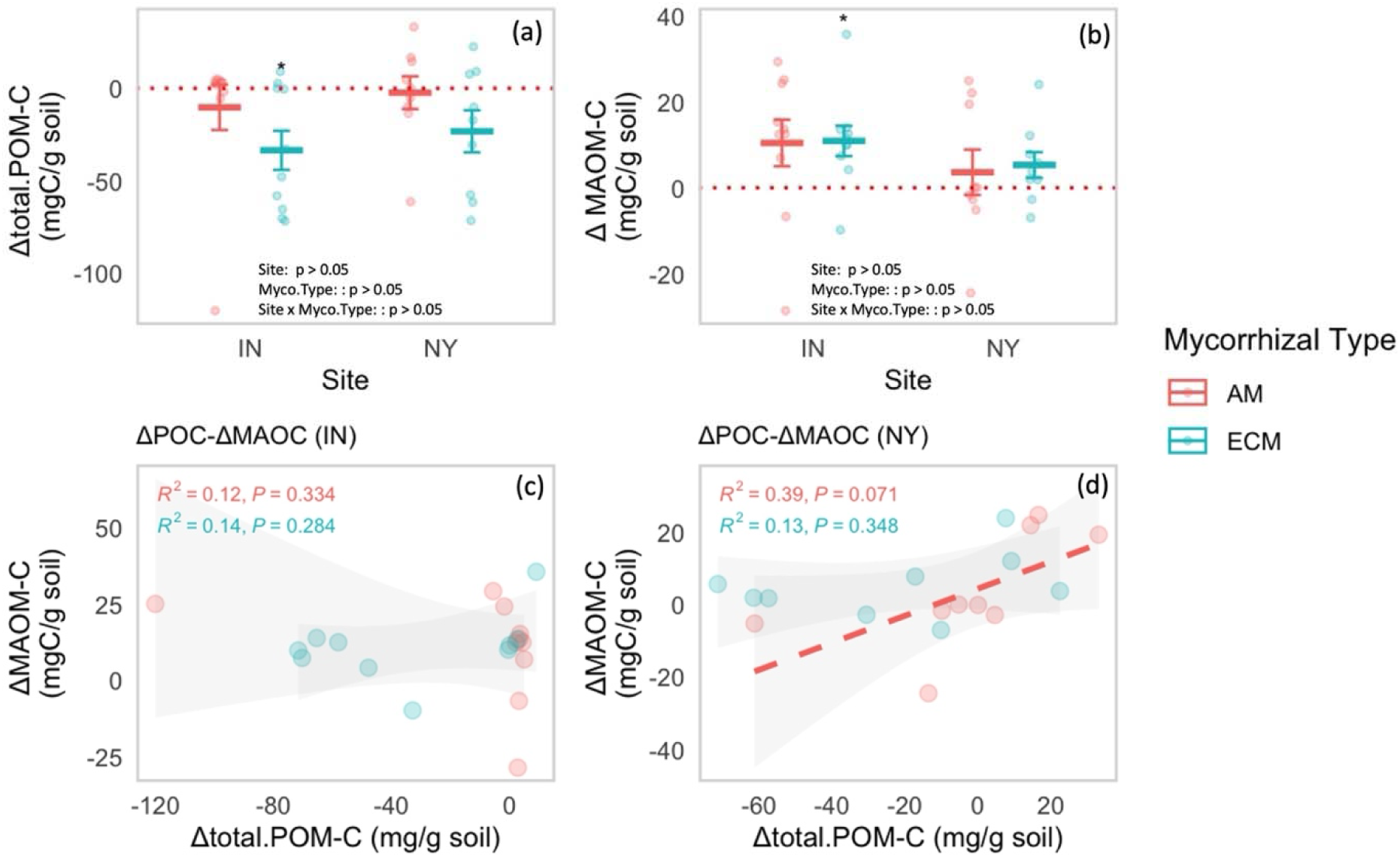
Changes in total particulate organic matter C (Δtotal POM-C) and mineral-associated organic matter C (ΔMAOM-C) and their relationships across sites and mycorrhizal types after 24 months of root ingrowth. Changes in (a) total particulate organic matter C (Δtotal POM-C; sum of changes in free and occluded POM-C) and (b) mineral-associated organic matter C (ΔMAOM-C) are expressed relative to initial soils (mg C g ¹ soil). Points represent individual observations, and crossbars indicate the mean ± SE for each Site × mycorrhizal type combination. Red dotted horizontal lines denote no change relative to initial soils (Δ = 0). Fixed effects of Site, mycorrhizal type, and their interaction were evaluated using linear models (lm), with model summaries shown within each panel. Asterisks indicate significant differences from zero within each Site × mycorrhizal type combination (P < 0.05). Panels (c) and (d) show relationships between Δtotal POM-C and ΔMAOM-C in Indiana (IN) and New York (NY), respectively. Lines represent fitted linear regressions for each mycorrhizal type, with shaded areas indicating 95% confidence intervals. Dashed lines indicate marginally significant relationships (P < 0.1), with model R^2^ and P-values shown within each panel. Colors indicate arbuscular mycorrhizal (AM) and ectomycorrhizal (EcM) plots throughout.

**Table 2).**
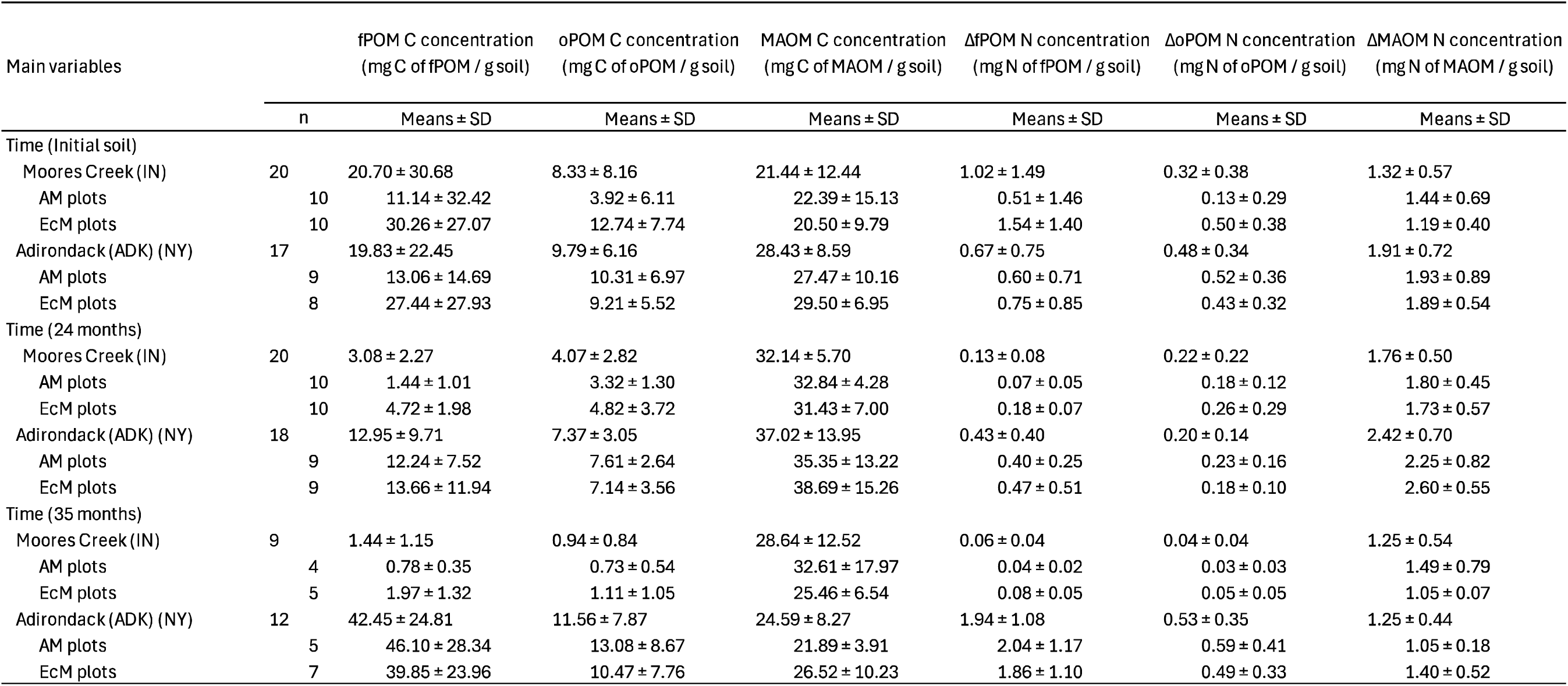
Means± SDs of soil organic matter C and N concentrations(mg g^-1^ soil) across particulate and mineral-associated fractions, Including free particulate organic matter (fPOM), occluded particulate organic matter (oPOM), and mineral-associated organic matter (MAOM)

| Main variables |  | fPOM C concentration<br>(mg C of fPOM / g soil) | oPOM C concentration<br>(mg C of oPOM / g soil) | MAOM C concentration<br>(mg C of MAOM / g soil) | ΔfPOM N concentration<br>(mg N of fPOM / g soil) | ΔoPOM N concentration<br>(mg N of oPOM / g soil) | ΔMAOM N concentration<br>(mg N of MAOM / g soil) |
|  | n | Means ± SD | Means ± SD | Means ± SD | Means ± SD | Means ± SD | Means ± SD |
|---|---|---|---|---|---|---|---|
| Time (Initial soil) |  |  |  |  |  |  |  |
| Moores Creek (IN) | 20 | 20.70 ± 30.68 | 8.33 ± 8.16 | 21.44 ± 12.44 | 1.02 ± 1.49 | 0.32 ± 0.38 | 1.32 ± 0.57 |
| AM plots | 10 | 11.14 ± 32.42 | 3.92 ± 6.11 | 22.39 ± 15.13 | 0.51 ± 1.46 | 0.13 ± 0.29 | 1.44 ± 0.69 |
| EcM plots | 10 | 30.26 ± 27.07 | 12.74 ± 7.74 | 20.50 ± 9.79 | 1.54 ± 1.40 | 0.50 ± 0.38 | 1.19 ± 0.40 |
| Adirondack (ADK) (NY) | 17 | 19.83 ± 22.45 | 9.79 ± 6.16 | 28.43 ± 8.59 | 0.67 ± 0.75 | 0.48 ± 0.34 | 1.91 ± 0.72 |
| AM plots | 9 | 13.06 ± 14.69 | 10.31 ± 6.97 | 27.47 ± 10.16 | 0.60 ± 0.71 | 0.52 ± 0.36 | 1.93 ± 0.89 |
| EcM plots | 8 | 27.44 ± 27.93 | 9.21 ± 5.52 | 29.50 ± 6.95 | 0.75 ± 0.85 | 0.43 ± 0.32 | 1.89 ± 0.54 |
| Time (24 months) |  |  |  |  |  |  |  |
| Moores Creek (IN) | 20 | 3.08 ± 2.27 | 4.07 ± 2.82 | 32.14 ± 5.70 | 0.13 ± 0.08 | 0.22 ± 0.22 | 1.76 ± 0.50 |
| AM plots | 10 | 1.44 ± 1.01 | 3.32 ± 1.30 | 32.84 ± 4.28 | 0.07 ± 0.05 | 0.18 ± 0.12 | 1.80 ± 0.45 |
| EcM plots | 10 | 4.72 ± 1.98 | 4.82 ± 3.72 | 31.43 ± 7.00 | 0.18 ± 0.07 | 0.26 ± 0.29 | 1.73 ± 0.57 |
| Adirondack (ADK) (NY) | 18 | 12.95 ± 9.71 | 7.37 ± 3.05 | 37.02 ± 13.95 | 0.43 ± 0.40 | 0.20 ± 0.14 | 2.42 ± 0.70 |
| AM plots | 9 | 12.24 ± 7.52 | 7.61 ± 2.64 | 35.35 ± 13.22 | 0.40 ± 0.25 | 0.23 ± 0.16 | 2.25 ± 0.82 |
| EcM plots | 9 | 13.66 ± 11.94 | 7.14 ± 3.56 | 38.69 ± 15.26 | 0.47 ± 0.51 | 0.18 ± 0.10 | 2.60 ± 0.55 |
| Time (35 months) |  |  |  |  |  |  |  |
| Moores Creek (IN) | 9 | 1.44 ± 1.15 | 0.94 ± 0.84 | 28.64 ± 12.52 | 0.06 ± 0.04 | 0.04 ± 0.04 | 1.25 ± 0.54 |
| AM plots | 4 | 0.78 ± 0.35 | 0.73 ± 0.54 | 32.61 ± 17.97 | 0.04 ± 0.02 | 0.03 ± 0.03 | 1.49 ± 0.79 |
| EcM plots | 5 | 1.97 ± 1.32 | 1.11 ± 1.05 | 25.46 ± 6.54 | 0.08 ± 0.05 | 0.05 ± 0.05 | 1.05 ± 0.07 |
| Adirondack (ADK) (NY) | 12 | 42.45 ± 24.81 | 11.56 ± 7.87 | 24.59 ± 8.27 | 1.94 ± 1.08 | 0.53 ± 0.35 | 1.25 ± 0.44 |
| AM plots | 5 | 46.10 ± 28.34 | 13.08 ± 8.67 | 21.89 ± 3.91 | 2.04 ± 1.17 | 0.59 ± 0.41 | 1.05 ± 0.18 |
| EcM plots | 7 | 39.85 ± 23.96 | 10.47 ± 7.76 | 26.52 ± 10.23 | 1.86 ± 1.10 | 0.49 ± 0.33 | 1.40 ± 0.52 |

The magnitude and direction of absorptive root density and SOM changes were similar between sites (Fig. 1). Both fPOM and oPOM generally declined following two years of root ingrowth, although significant decreases were observed only in EcM-dominated plots at IN. EcM plots at IN, in particular, exhibited significant decreases in both fPOM-C (−83%) and oPOM-C (−58%), contributing to the greater overall decline in POM-C observed in these plots (p <0.05; Fig. 1a; 1b; Table 2, S6). Changes in POM-N varied among sites and mycorrhizal types, with significant declines in oPOM-N observed in both AM and EcM plots at NY (Fig. S3). In contrast to the POM fractions, MAOM-C generally increased across both sites and tree mycorrhozal types (Fig. 1c). Increases in MAOM-C were generally greater in IN, though this may reflect the fact that NY contained a significantly larger initial MAOM-C pool prior to root ingrowth (Fig. S4). Increases in MAOM-N differed slightly from MAOM-C, as only ECM plots in NY increased in response to root ingrowth (Fig S4).

### Relationships between root traits and SOM-C dynamics after 24 months

Principal component analysis (PCA) of six root morphological traits and root density identified two orthogonal axes of root trait variation after two years of root ingrowth. The first principal component (PC1), primarily associated with specific root area (SRA) and specific root length (SRL), explained 48% of the total variation, while the second component (PC2), primarily associated with branching intensity (BI), accounted for an additional 28% (Fig. S5c,d; Table S5). Across all 24-month samples (n = 38), neither ΔPOM-C nor ΔMAOM-C were significantly associated with multivariate root trait organization (Fig. S5c). PERMANOVA further revealed significant differences in root trait organization between sites (R^2^ = 0.159, P < 0.001), mycorrhizal types (R^2^ = 0.076, P < 0.001), and sampling times (R^2^ = 0.117, P < 0.001), but no significant interaction between mycorrhizal type and sampling time (R^2^ = 0.024, P = 0.056). Multivariate dispersion did not differ significantly among sites, mycorrhizal types, sampling times, or their combinations (all P > 0.18), indicating that differences in root trait organizations were not accompanied by detectable differences in within-group dispersion.

Given the differences in multivariate root trait organization between mycorrhizal types, we conducted separate PCAs for AM- and EcM-dominated plots (n = 19 per group; Fig. 3; Table S5). In EcM plots, ΔMAOM-C was negatively associated with PC1 (Fig. 4b), which reflected most variation in root morphological traits, particularly SRL, SRA and root tissue density (RTD). Accordingly, greater MAOM-C accumulation was associated with lower SRA and higher RTD (Fig. 4a,b). In contrast, although ΔMAOM-C tended to decline along PC1 in AM plots, this relationship was not statistically significant. No individual root traits were significantly linked to ΔPOM-C or ΔMAOM-C in AM plots.

**Fig. 3).**
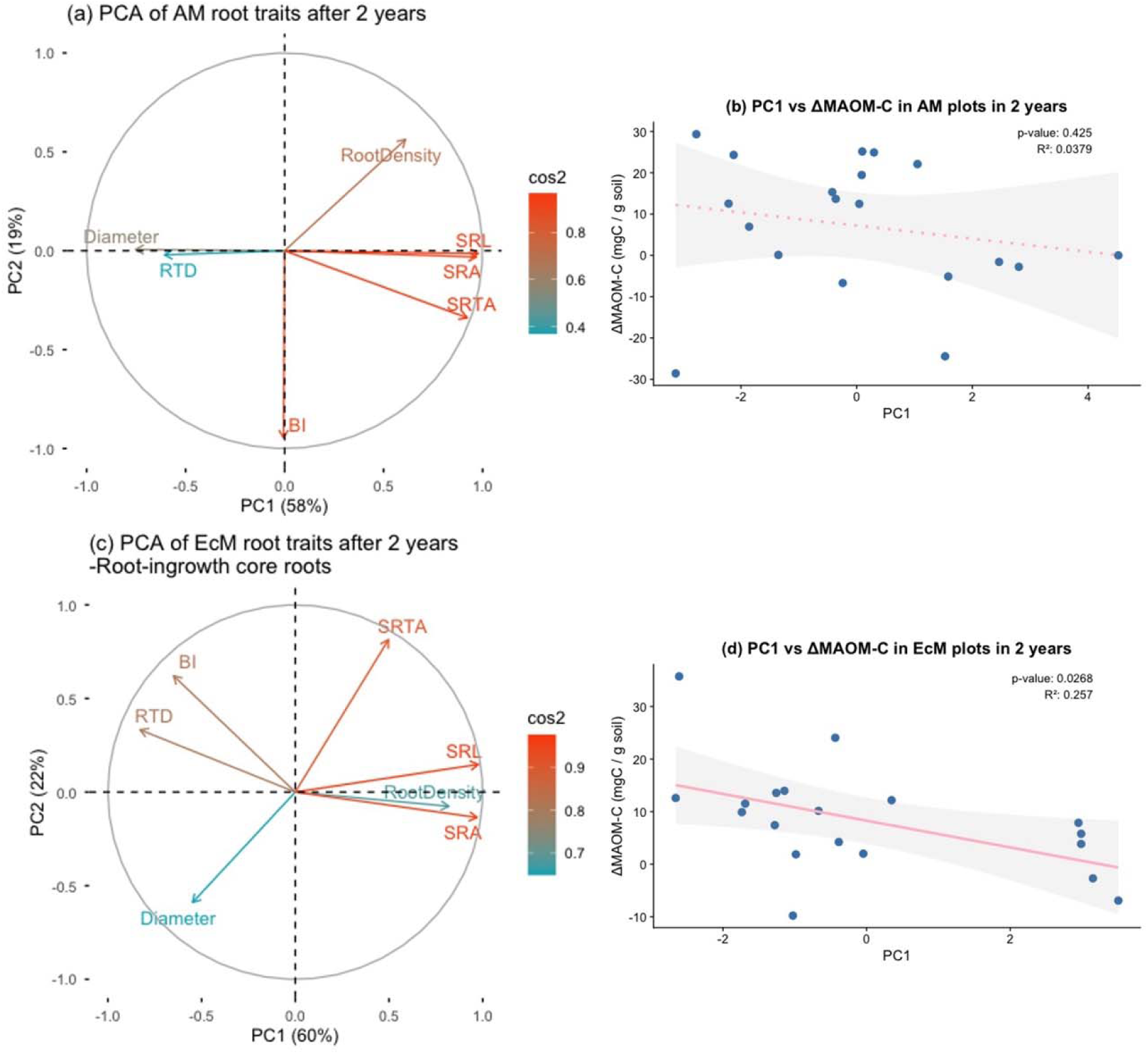
Root trait organization and its relationships with changes in mineral-associated organic matter C (MAOM-C) in arbuscular mycorrhizal (AM)-and ectomycorrhizal (EcM)-dominated forests after two years of root ingrowth. (a, c) Principal component analyses (PCA) of seven fine-root traits in AM-(n=19)(a) and EcM-dominated forests (n=19)(c). The first two principal components explained 77% and 82% of total trait variation in AM and EcM forests, respectively. Arrows indicate trait loadings, with colors representing the quality of representation (cos^2^) on the first two principal components. (b, d) Relationships between PC1 scores and changes in MAOM-C concentrations (ΔMAOM-C) relative to initial soils in AM (b) and EcM forests (d). PC1 was negatively associated with ΔMAOM-C in EcM forests (R^2^ = 0.257, P = 0.027), whereas no significant relationship was observed in AM forests (R^2^ = 0.038, P = 0.425). Points represent individual observations, solid lines indicate linear regressions, and shaded areas represent 95% confidence intervals. Separate PCAs were conducted for each mycorrhizal type to characterize mycorrhizal-specific root trait organization.

**Fig. 4).**
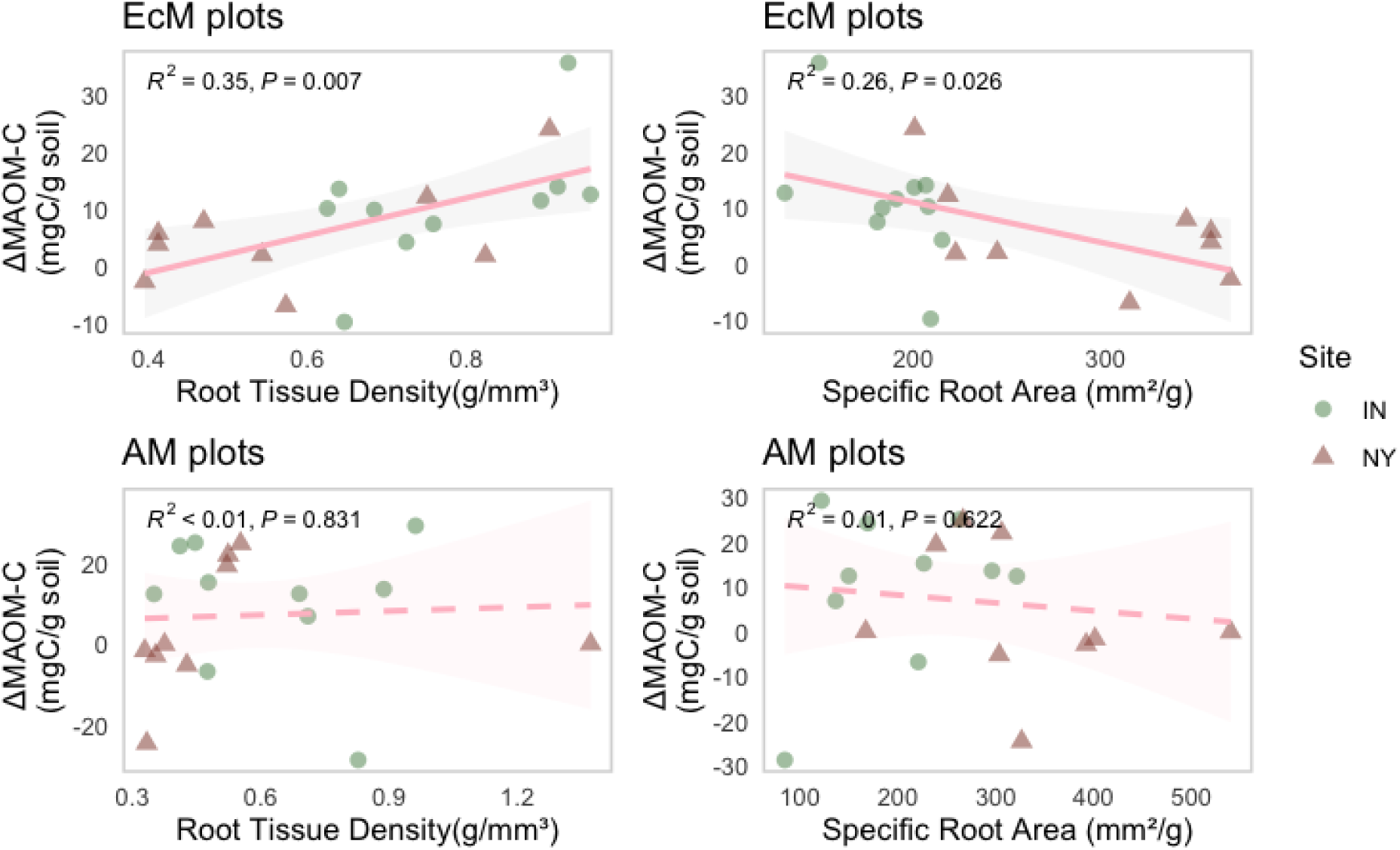
Relationships between changes in mineral-associated organic matter C (ΔMAOM-C) and root tissue density (RTD; g/mm³) and specific root area (SRA; mm^2^/g) in EcM and AM plots across sites. RTD and SRA were selected for further examination because they were the two highest-ranked individual predictors of ΔMAOM-C in the redundancy analysis (RDA) model-selection procedure conducted for EcM plots (adjusted R^2^ = 0.313 and 0.217, respectively; Fig. S00). However, only RTD showed a significant relationship with ΔMAOM-C, and including SRA did not improve the RTD-only model. Upper and lower panels represent EcM- and AM-dominated plots, respectively. Points represent individual observations, with point shape and color indicating site (IN and NY). Lines represent fitted linear regressions (solid for significant relationships and dashed for nonsignificant relationships), with shaded areas indicating 95% confidence intervals. Model R^2^ and P-values are shown within each panel.

Further RDA analyses of EcM plots identified RTD as the strongest individual predictor of ΔMAOM-C (Adj. R^2^ = 0.31), followed by SRA (Adj. R^2^ = 0.22) and root density (Adj. R^2^ = 0.16). Consistent with these patterns, ΔMAOM-C was positively associated with RTD (R^2^ = 0.35, P = 0.007) and negatively associated with SRA (R^2^ = 0.26, P = 0.03; Fig. 4a,b). Although a model including both RTD and SRA was significant (P = 0.025), SRA did not explain additional variation after accounting for RTD (P = 0.955), suggesting that RTD accounted for most of the variation explained by these two traits.

### Effects of nutrient addition on root–SOM dynamics in two years

Bi-monthly nutrient additions over two growing seasons had no significant effects on absorptive root density (Fig. S18) but in some cases, affected the relationships (slopes) between absorptive root density and SOM (Fig. 5; S7). As such, fertilization had no effects on the relaionship between root density and POM-C change but did alter the relationship between MAOM-C and root-density (Fig. 5; S7). In control cores at the NY site, ΔMAOM-C was negatively related to absorptive root density (p = 0.024; Fig. 3b), whereas this relationship was not significant under nutrient addition and instead tended to be positive (p = 0.089; Fig. 3a). A similar shift in the direction of the root–MAOM-C relationship was apparent at the IN site, although neither relationship was significant. The relationships between root density and ΔMAOM-C were unaffected by tree-mycorrhizal type (Fig. S7).

**Fig. 5).**
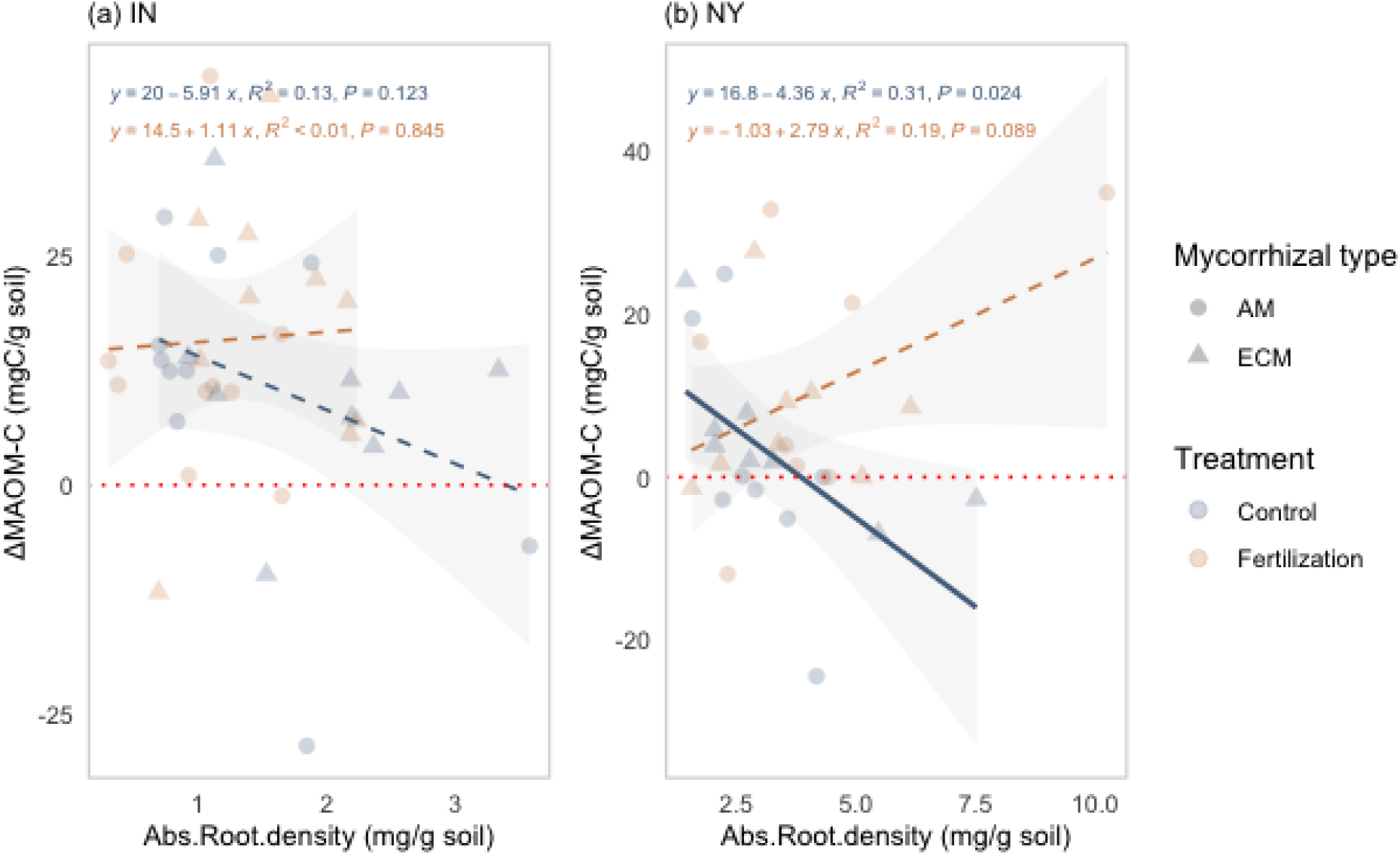
Effects of fertilization on relationships between absorptive root ingrowth density and mineral-associated organic matter C formation (ΔMAOM-C) at (a) IN and (b) NY. ΔMAOM-C represents the change in MAOM-C concentration relative to initial soils (mg C g ¹ soil). Points represent individual observations, with point shape indicating mycorrhizal type (AM and ECM) and color indicating treatment (control and fertilization). Lines represent treatment-specific linear regressions, with shaded areas indicating 95% confidence intervals. Differences in root–MAOM relationships between control and fertilized cores were evaluated by testing the absorptive root density × treatment interaction. Model R^2^ and P-values for treatment-specific relationships are shown within each panel. Note) The effects of fertilization on relationships between absorptive root ingrowth density and the total ΔPOM-C at IN and NY were not significant (Fig. S00)

Nutrient addition had no or little detectable effect on ΔPOM-C, with paired treatment differences remaining nonsignificant at both IN and NY (Fig. S11). However, consistent with changes in root–MAOM relationships, nutrient addition generally enhanced MAOM-C accumulation, with the magnitude of this response varying between sites (Fig. S10). For instance, paired comparisons revealed greater MAOM-C accumulation under nutrient addition at the NY site, where higher mean soil C:N (20.7 vs. 19.2 at IN; Fig. S8) putatively suggested greater N limitation. Nutrient addition significantly increased ΔMAOM-C in both AM-(P = 0.050; Fig. S9) and EcM-dominated plots (P = 0.039; Fig. 4). At IN, ΔMAOM-C also tended to increase under nutrient addition, although paired differences were not significant for either AM or EcM plots (Fig. 4). Overall, the SOM-C response to nutrient addition was revealed through changes in MAOM-C rather than POM-C, and the responses were most pronounced in EcM plots in NY (Fig. S9).

### Changes in root-induced SOM dynamics over time

Root-induced changes in SOM pools varied over time, among SOM fractions, and between sites (Fig. 5). In general, changes in SOM fractions from 24 months to 35 months were greatest in IN. Over the first 24 months of ingrowth, roots induced reductions in fPOM-C and oPOM-C (p < 0.05; Fig. 5a); after 35 months, these effects persisted for fPOM-C and the increased in magnitude (i.e., greater losses) for oPOM-C (p < 0.05; Fig. 6a). In contrast, MAOM-C in IN remained elevated relative to initial soils over the three-year period (p < 0.05; Fig. 6a). In NY, we observed no significant changes in fPOM-C, oPOM-C or MAOM-C between 24 and 35 months (Fig. 5a). Thus, the transition from two to three years of root ingrowth was characterized by (1) contrasting SOM-C trajectories between sites, (2) continued reduction in POM-C at IN but no changes in POM-C at NY, and (3) diminishing MAOM-C gains at NY. Changes in SOM-N showed fraction- and site-dependent temporal patterns, although their magnitudes were considerably smaller than those observed for SOM-C (Fig. 5b). Changes in fPOM-N were generally small, whereas oPOM-N tended to decline relative to initial soils, particularly after 24 months (p > 0.05; Fig. 6b). MAOM-N responses were more variable among sites and through time. In IN, positive MAOM-N changes after 24 months diminished or became negative by 35 months whereas in NY, MAOM-N, which had changed little over 24 months, declined by 35 months. Taken together, our results suggest that root-induced redistribution of C and N among POM and MAOM pools was dynamic over the three-year period.

**Fig. 6a).**
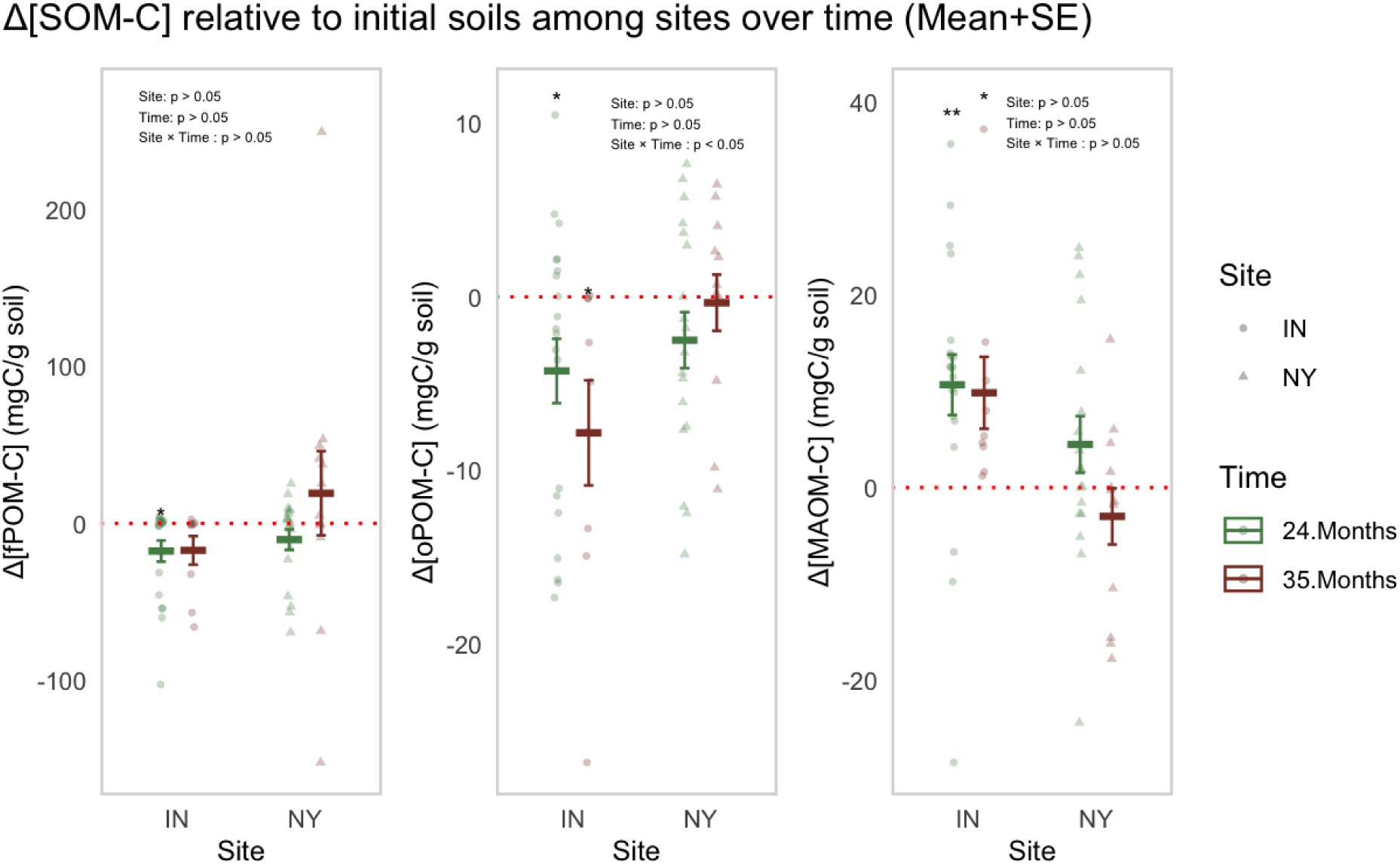
The effects of Site, Time, and their interaction on changes in soil organic matter C concentrations relative to initial soils (ΔSOM-C; mg C g ¹ soil) in the free particulate organic matter (fPOM), occluded particulate organic matter (oPOM), and mineral-associated organic matter (MAOM) fractions. Points represent individual observations, and crossbars indicate the mean ± SE for each Site × Time combination. Red dotted horizontal lines denote no change relative to initial soil (Δ = 0). Fixed effects of Site, Time, and their interaction were evaluated using linear mixed-effects models with Plot included as a random effect. Asterisks indicate whether ΔSOM-C differed significantly from zero within each Site × Time combination (p < 0.05, p < 0.01), while model summaries are shown within each panel.

**Fig. 6b).**
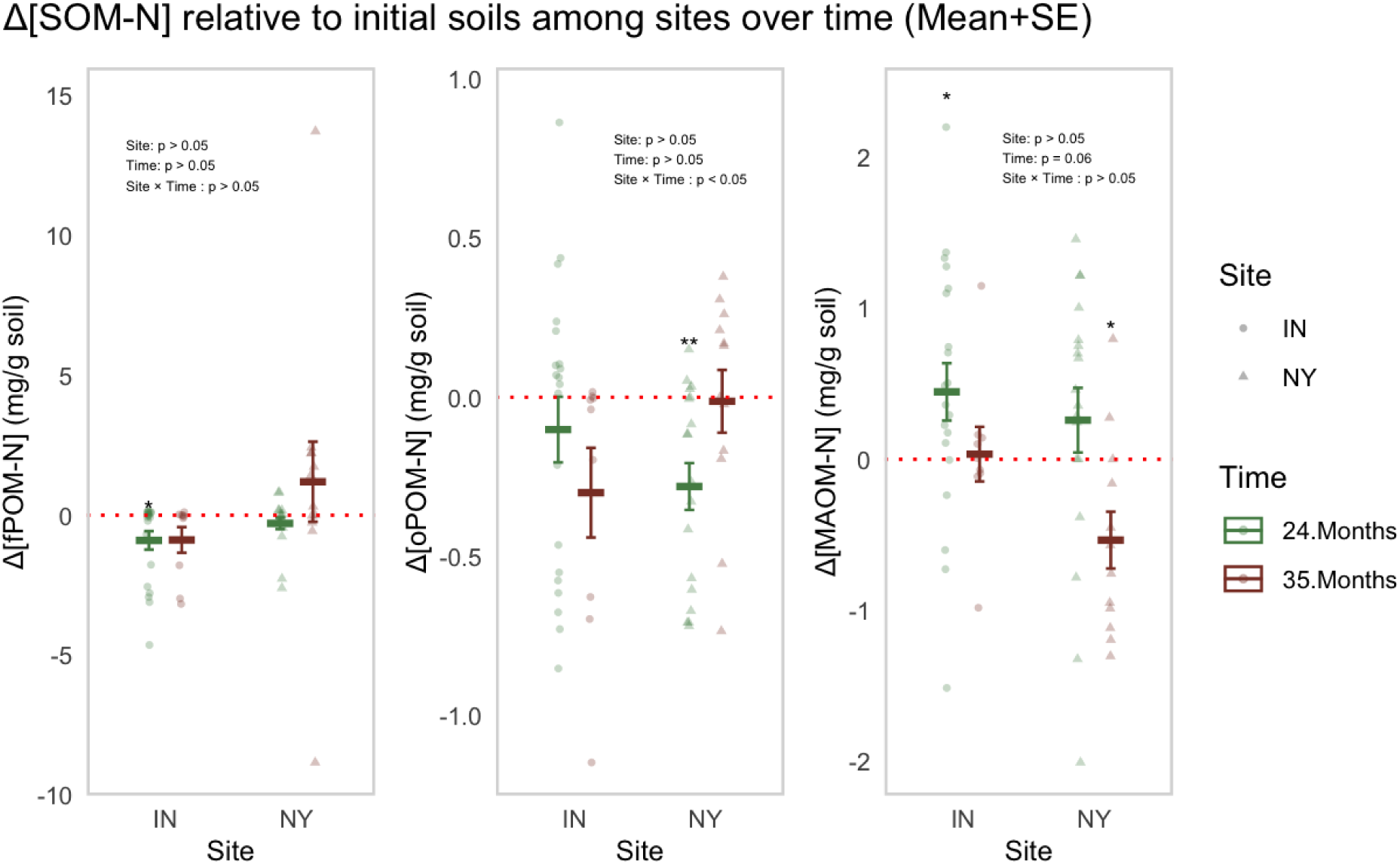
The effects of Site, Time, and their interaction on changes in soil organic matter N concentrations relative to initial soils (ΔSOM-N; mg N g ¹ soil) in the free particulate organic matter (fPOM), occluded particulate organic matter (oPOM), and mineral-associated organic matter (MAOM) fractions. Points represent individual observations, and crossbars indicate the mean ± SE for each Site × Time combination. Red dotted horizontal lines denote no change relative to initial soil (Δ = 0). Fixed effects of Site, Time, and their interaction were evaluated using linear mixed-effects models with Plot included as a random effect. Asterisks indicate whether ΔSOM-C differed significantly from zero within each Site × Time combination (p < 0.05, p < 0.01), while model summaries are shown within each panel.

Variation among forest locations within the NY site further revealed that site-level patterns masked substantial differences in root-SOM dynamics (Figs. S10). The positive MAOM-C response observed at NY after 24 months (Fig. S10a) was driven largely by Lake George Wild Forest (DL), where MAOM-C also increased significantly relative to initial soils. This location also showed a significant positive change in oPOM-C after 24 months (p < 0.01; Fig. S10a). By 35 months, however, both responses had diminished toward initial-soil levels (Fig. S10a). A similar positive MAOM-C response after 24 months occurred at IN, but, unlike DL, MAOM-C at IN remained elevated after 35 months (Fig. S10a). In contrast, the other two NY forest locations showed little evidence of sustained MAOM-C accumulation: MAOM-C tended to decline at Shingle Shanty Nature Preserve (SS), whereas changes at Huntington Wildlife Forest (HF) remained relatively small.

The contrasting SOM trajectories among the three NY forest locations appeared to be linked to pronounced differences in their environmental and edaphic characteristics. DL had the highest climate decomposition index (CDI) among the NY locations (0.43 vs. approximately 0.38 at SS and HF; Table 1), higher soil pH (5.1 vs. 3.7–3.9; Fig S16), and lower mean soil C:N (19.2 vs. 20.7 at NY; ; Table 1) than the other NY locations (Eagar et al., 2022). SS and HF, in contrast, were characterized by lower CDI and more acidic, C-rich soils (Eagar et al., 2022). These stand-level differences accompanied divergent SOM responses: DL exhibited relatively strong MAOM-C accumulation after 24 months followed by a return toward initial levels, whereas SS and HF showed weaker or negative MAOM-C responses (Fig. S14a). Thus, variation among forest locations accounted for an important component of the temporal heterogeneity in root-induced SOM dynamics observed at the broader NY site.

## Discussion

While roots are well known to influence SOM dynamics (Rasse et al., 2005; Bardgett et al., in review), studies of root growth and SOM change from the same experimental unit remain relatively rare. Here, we investigated how root affect SOM fractions in forest plots dominated by arbuscular mycorrhizal (AM) and ectomycorrhizal (EcM) trees using root ingrowth cores. We tested three hypotheses: H1) root-induced changes in SOM fractions differ between tree-mycorrhizal types, with EcM roots promoting POM losses and AM roots enhancing MAOM gains, H2) increases in nutrient availability reduce POM loss and enhance MAOM formation, with the greatest changes occurring in more N-limited environments, and H3) root-induced changes in POM and MAOM fluctuate over time, as initial gains in SOM are offset by SOM losses. Overall, we some support for each hypothesis. After two years, root ingrowth was generally associated with losses of POM-C and gains in MAOM-C, with EcM-dominated plots showing particularly greater POM-C losses and significant MAOM-C accumulation. Nutrient addition altered this relationship and enhanced MAOM-C accumulation, particularly at the more N-limited EcM dominated plots at NY, while resulting in comparatively little or no change in POM-C. Importantly, the initial increase in MAOM-C was not persistent throughout 35 months of root ingrowth. By 35 months at NY, MAOM-C gains had diminished toward initial-soil levels, while MAOM-C gains at IN remained elevated. These findings collectively suggest that root effects on SOM cannot be predicted from new root inputs alone, but instead emerge from interactions among mycorrhizal roots’ strategies, nutrient availability, environmental controls on decomposition and time.

### Two years of root ingrowth redistributed SOM-C among fractions

The degree to which roots influence SOM stocks in forests is unclear, as roots can accelerate losses via priming (Dijkstra & Cheng, 2007), slow losses by enhancing mineral stabilization (Heckman et al., 2023; Lang et al., 2026), or have little to no net effects if SOM acceleration/stabilization processes are offsetting (Malhotra et al., 2025). In this study, two years of root ingrowth generally resulted in POM-C losses and MAOM-C gains, demonstrating that root effects lead to redistribution of C among SOM fractions rather than uniform accumulation of soil C. Across sites, these responses differed with plot mycorrhizal dominance: EcM plots showed significant MAOM-C accumulation but also substantially greater POM-C losses, ultimately resulting in a net total SOM-C loss (Fig. S1). In contrast, total SOM-C remained near initial levels in AM plots and tended to be greater than in EcM plots (Fig. S1). These patterns are consistent with previous studies showing that greater EcM root proliferation does not necessarily result in greater soil C retention (Keller et al., 2021; Malhortra et al., 2025; Lang et al., 2026). For instance, Keller et al. (2021) used ingrowth cores packed with isotopically-distinct SOM-C and found that greater root growth in EcM-dominated plots did not coincide with greater root-derived C inputs to soil or changes in SOM fractions. In contrast, AM-dominated stands retained a larger fraction of root-derived C in MAOM. The relationship between fine root biomass (or C inputs) and SOM-C (or MAOM-C) in forests remains ambiguous, as some studies have reported no relationship (Malhortra et al., 2025) whereas others have reported positive relationships (Heckman et al. 2023; Muratore et al. 2026; Lang et al. 2026). Whether this paradox relates to how AM and EcM roots modify POM and MAOM is unclear. Root-induced MAOM formation can depend on tree-mycorrhizal type differences, as EcM-associated roots may stimulate microbial processing and promote MAOM formation while simultaneously mining new and existing particulate SOM (Dijkstra & Cheng, 2007; Liang et al., 2018). Such simultaneous MOAM formation and POM loss processes could explain why greater EcM absorptive root density in our study coincided with significant MAOM-C accumulation (Fig. 1) but not greater total SOM-C retention (Fig. S1).

Beyond mycorrhizal dominance, site-specific environmental conditions appeared to regulate the magnitude of root-associated POM-C losses and MAOM-C gains. We found that POM-C losses were more pronounced at IN, where a higher climatic decomposition index (CDI), higher soil pH, and narrower soil C:N may have provided conditions more favorable for SOM decomposition than at NY. EcM plots at IN exhibited particularly strong fPOM-C and oPOM-C losses, potentially reflecting that the magnitude of root-induced decomposition of POM depends on the abundance of EcM roots that have a capability to degrade recalcitrant SOM using extracellular enzyme (Comas et al., 2014; Courty et al., 2010). Greater POM losses at IN were also in line with greater MAOM-C gains, despite the larger initial MAOM-C pool at NY (Fig. S4). These findings are consistent with our general understanding on environmental constraints for SOM formation, in which rhizosphere conditions favorable for decomposition not only accelerate the processing of POM and root-inputs (Dijkstra et al., 2021; Dijkstra & Cheng, 2007), but also simultaneously promote MAOM-C formation by microbially processed organic matter compounds (Liang et al., 2017; Sokol et al., 2019; Sokol & Bradford, 2019). Therefore, the concurrent greater POM-C losses and MAOM-C gains at IN may reflect greater SOM transformation, whereby enhanced processing of organic matters can subsequently contribute to MAOM formation.

Interestingly, greater absorptive root ingrowth did not correspond to greater MAOM-C accumulation. Instead, absorptive root density was negatively associated with MAOM-C gain across sites, although this relationship was driven primarily by the NY sites and was not detectable at IN (Fig. S6). These results pose a new question on whether greater root biomass in forests generally results in greater MAOM formation (Heckman et al., 2023; Malhortra et al., 2025) and how the relationships differ among forests. Rather, net changes in MAOM in our results represent the balance between root-induced formation and loss processes (Dijkstra & Cheng, 2007). As such, greater root proliferation can increase soluble labile C inputs such as root exudates (Oh et al., 2026) and microbial products that contribute to MAOM formation (Jackson et al., 2017; Rasse et al., 2005), while simultaneously stimulating microbial activity, nutrient acquisition, or rhizosphere chemical conditions capable of mobilizing existing SOM (Cheng et al., 2014; Finzi et al., 2015; Muratore et al., 2026). The absence of root-MAOM relationship at IN, which was also observed by previous studies (Keller et al., 2021; Malhortra et al., 2025; Lang et al., 2026), further suggests that this balance likely depends on environmental context rather than representing a general consequence of root proliferation.

Importantly, variation in root traits was found to associate with MAOM-C dynamics in EcM-dominated plots, but these relationships were largely absent in AM-dominated plots, suggesting that tree-mycorrhizal dominance may influence how root traits affect SOM dynamics. Indeed, two years of root ingrowth induced root traits that differed from those observed in initial soils (Fig. S5a,c), with contrasting relationships between trait organization and MAOM-C changes across mycorrhizal types. In EcM plots, the first principal component (PC1), associated with greater specific root length (SRL), specific root area (SRA), and root density, was negatively related to ΔMAOM-C, whereas AM plots demonstrated no detectable relationship (Fig. 3d). Consistent with this pattern, SRA was negatively associated with ΔMAOM-C in EcM plots (Fig. 3,4), suggesting that more acquisitive root morphologies may link to a reduced net MAOM-C accumulation. Conversely, RTD, placed on the conservative end of the root economics spectrum (Freschet et al., 2021; Weigelt et al., 2021), was positively related to ΔMAOM-C in EcM plots (Fig. 4). This association may reflect differences in root structural investment, tissue longevity, or mycorrhizal contributions to SOM formation (Chen et al., 2025; Hawkins et al., 2023). However, whether higher RTD corresponds to greater EcM colonization or fungal biomass remains uncertain in our study, making it difficult to distinguish the contributions of root tissue properties from those of associated fungi. In contrast, the weaker coordination among root morphological traits and their unclear associations with ΔMAOM-C in AM plots may indicate that morphological traits alone may be insufficient to predict their effects on SOM dynamics. AM roots, for example, may instead influence MAOM-C formation via processes less directly attributable to root morphology, such as exudation-mediated microbial processes and nutrient mobilization (Lv et al., 2026). Nonetheless, these potential mechanisms should remain speculative given that neither root-derived C inputs nor fungal contributions to MAOM formation were directly quantified. Further field-based studies integrating root traits, rhizodeposition, mycorrhizal fungal activity, and SOM turnover will be central to determine how these processes jointly regulate MAOM-C dynamics across mycorrhizal types.

### Nutrient modulation of root-SOM relationships

Our bi-monthly nutrient addition experiment revealed that root effects on SOM dynamics depended on site-specific nutrient availability and potentially, environmental controls over decomposition. In line with our hypothesis (H2) that fertilization effects would be greater where nutrient limitation was stronger, nutrient addition enhanced MAOM-C accumulation, with the strongest responses occurring at NY, where lower pH, higher soil C:N, more recalcitrant litter inputs, and lower climate decomposition potential (Table 1) are consistent with slower nutrient turnover. In contrast to the IN site, these changes were accompanied by a shift in the root–MAOM-C relationship at NY, from a significant negative relationship under ambient conditions to a nonsignificant, but positive, relationship under nutrient addition (Fig. 5). These responses suggest that nutrient availability may regulate not only the magnitude of MAOM formation but also how increasing root inputs influence the balance between SOM transformation and MAOM accumulation (Dijkstra et al., 2021; Dijkstra & Cheng, 2007).

One potential explanation for the stronger fertilization response at NY is that nutrient addition altered microbial and mycorrhizal nutrient-acquisition activities that mediate the fate of root-derived and existing soil C. Previous work conducted across the same NY forests demonstrated that fungal community composition was related to tree-mycorrhizal dominance and environmental factors (Eagar et al., 2023). Cortese et al. (2026) further reported that ECM fungal functional composition across these forests was strongly associated with soil conditions, including pH, C:N, and N availability. ECM roots’ strategies characterized by greater investment in extramatrical nutrient foraging may be advantageous under more N-limited conditions but require substantial host C investment (Chen et al., 2025). Thus, alleviation of nutrient limitation could reduce investment in belowground nutrient-foraging and consequently alter microbial processing of SOM. In line with such a shift in belowground foraging, EcM root branching intensity (BI), which is known to be closely linked to roots’ C investment to mycorrhizal partners (Brundrett et al., 1990; Comas & Eissenstat, 2009), declined significantly under nutrient addition at NY (p < 0.05; Fig. S20b), and these reduced BI were coincided with greater MAOM-C accumulation (Fig. S20b). Although decreases in root branching do not directly indicate reduced C allocation to EcM fungi, these concurrent responses suggest that alleviating nutrient limitation may alter belowground C allocation and nutrient-acquisition strategies of mycorrhizal roots, potentially changing the balance between C used for nutrient acquisition and C processed into pathways contributing to MAOM formation.

### Root-induced SOM dynamics over time

Our results from three years of root ingrowth revealed that MAOM accumulation was dynamic rather than continuously increasing with root ingrowth. We found increasing trends in MAOM-C across sites after 24 months, while the C gains had diminished toward initial-soil levels by 35 months (Fig. 6a; Fig. S4). In particular, the dynamics of C gains from 24 months to 35 months differed between the two sites. At IN, MAOM-C remained elevated while POM-C continued to decline, but at NY, MAOM-C gains diminished by 35 months while POM-C tended to recover and rather increase. The decline in MAOM-C gains between 24 and 35 months at NY suggests that root-induced MAOM accumulation likely represents a dynamic balance between SOM formation and loss rather than a continuously accumulating C sink (Dijkstra & Cheng, 2007; Liang et al., 2018). Although our measurements on SOM C and N concentration (Fig. S4) cannot distinguish declining MAOM formation from increasing loss of existing MAOM, the observed temporal response on root ingrowth demonstrates that MAOM accumulated during the initial period of root colonization was not uniformly maintained through time (between two to three years). Studies on how and the extent of roots’ effects on SOM for longer than three years are rare, and we still have a limited capability to cross-compare the results from different sites, calling for longer-term root ingrowth core studies across ecosystems.

The contrasting POM and MAOM dynamics between 24 and 35 months also suggest that the fate of root-derived SOM inputs changed differently between ecosystems. At IN, continued POM-C losses coincided with continuous MAOM-C gains, possibly sustaining transformation of POM to MAOM under environmental conditions favorable for decomposition (i.e., higher CDI: 0.53 vs. 0.40; Table 1) and subsequent retention of MAOM-C pools. In contrast, the increases in POM-C at NY coincided with the net reduction of initial MAOM-C gain, suggesting that with increasing root ingrowth, net C accumulation shifted toward POM fractions rather than mineral-associated pools. These site-specific dynamics indicate that root effects on SOM persistence depend not only on the magnitude of belowground inputs but also on how those inputs and existing SOM are processed and retained through time.

Resolving the two sites into four individual forest locations provides further evidence that MAOM dynamics were not spatially uniform. For example, the apparent MAOM-C gains at NY after 24 months were driven primarily by Lake George Wild Forest (DL), whereas Shingle Shanty Nature Preserve (SS) tended to lose MAOM-C and Huntington Wildlife Forest (HF) showed relatively little change. One reason for the greater MAOM-C gains at DL could be narrower soil C:N (16.4 vs. 22.8), higher pH (5.1 vs. 3.8) and CDI (0.426 vs. 0.385; warm/dry vs. cool/wet) than other two forest locations (SS and HF)(Eagar et al., 2023), all of which allows for expecting a higher microbial decomposition activities that lead to greater MAOM formation. In addition, it is worth noting that even the pronounced initial MAOM-C gains at DL diminished back to the initial level by 35 months (Fig. S10), reflecting a prevailing priming of MAOM over MAOM formation. These differences among locations suggest that the magnitude of root-induced MAOM formation and its persistence may depend strongly on environmental controls over SOM transformation.

Together, the reversal of MAOM responses to root ingrowth between 24 and 35 months highlights the importance of experimental duration when evaluating root effects on soil C persistence. For example, if responses were evaluated only after two years, the observed MAOM gains could have been interpreted as persistent OM stabilization, whereas our third-year results revealed that much of this response might be transient, particularly when nutrient availability is low and environmental controls for decomposition are relatively unfavorable. Therefore, longer-term observations (> 3 years) may be needed to determine whether root-induced MAOM gains represent sustained changes in soil C storage or short-term shifts in the balance between SOM formation and loss. Such studies would be especially useful across ecosystems in transition (e.g., persistent shrub invasion, new tree recruitments in grasslands, tree and shrub establishment in permafrost regions), where shifting plant communities may lead to unpredictable patterns of SOM gains and losses.

## Considerations and limitations

While our three-year field experiment identified important environmental factors associated with root effects on SOM dynamics, several limitations constrain our ability to determine the mechanisms underlying the balance between SOM formation and loss in the rhizosphere. First, although nutrient addition significantly increased MAOM-C at NY, where nutrient limitation was presumably greater, most measured root traits did not respond significantly to inorganic nutrient addition. Whether roots responded physiologically – exuding more or different types of C – was unmeasured abnd this could have affected the root-SOM relationships. Second, while the observed changes C and N in SOM relative to plot-specific initial soils allowed us to evaluate net changes in POM and MAOM, we were unable to distinguish the degree to which roots vs mycorrhiozal fungi vs free-living microbes contributed to those changes. Third, inherent differences in soil mineralogy between sites may have contributed to the contrasting MAOM changes over time. The NY sites contained acidic Spodosols, whereas soils at IN were predominantly Inceptisols (Table 1). Variations in O horizon thickness, pH, soil C:N, and climate can all control SOM stabilization and future studies should aim to separate the contributions of each effect. Lastly, the increase from ambient pH of roughly 5.5/4.2 to 7 in the IN cores is large enough to consider alternative explanations on the observed increases and persistence of MAOM-C at IN over 35 months. These limitations highlight the needs for future root ingrowth studies that can differentiate sources (old vs. new), processes (rhizosphere priming vs. formation), and resulting persistence using a combination of mechanistic isotopic tracing and field observations. Such approaches could also determine how living and dead roots differentially affect SOM fractions, and how and to what extent labile root C inputs can alter the balance between C and N retention through changes in microbial community composition and activity.

## Conclusion

Our findings reveal that root ingrowth can simultaneously promote POM-C losses and MAOM-C gains, with the balance between these responses varying across mycorrhizal types and environmental conditions. Greater root proliferation did not necessarily translate into greater soil C retention leading to contrasting SOM dynamics between IN and NY site. Nutrient addition that expected to stimulate microbial processing of SOM enhanced MAOM-C accumulation, particularly at the more N-limited NY site. Importantly, MAOM-C gains observed after two years were not consistently maintained after three years, highlighting the dynamic nature of root–SOM interactions. These findings collectively suggest that predicting root contributions to soil C persistence requires consideration of both SOM formation and loss processes, their environmental controls such as climate, edaphic factors, nutrient availability, and how their relative importance changes over time.

