## Supporting_Information for "Tree roots drive POM losses and transient MAOM gains across AM and ECM stands"

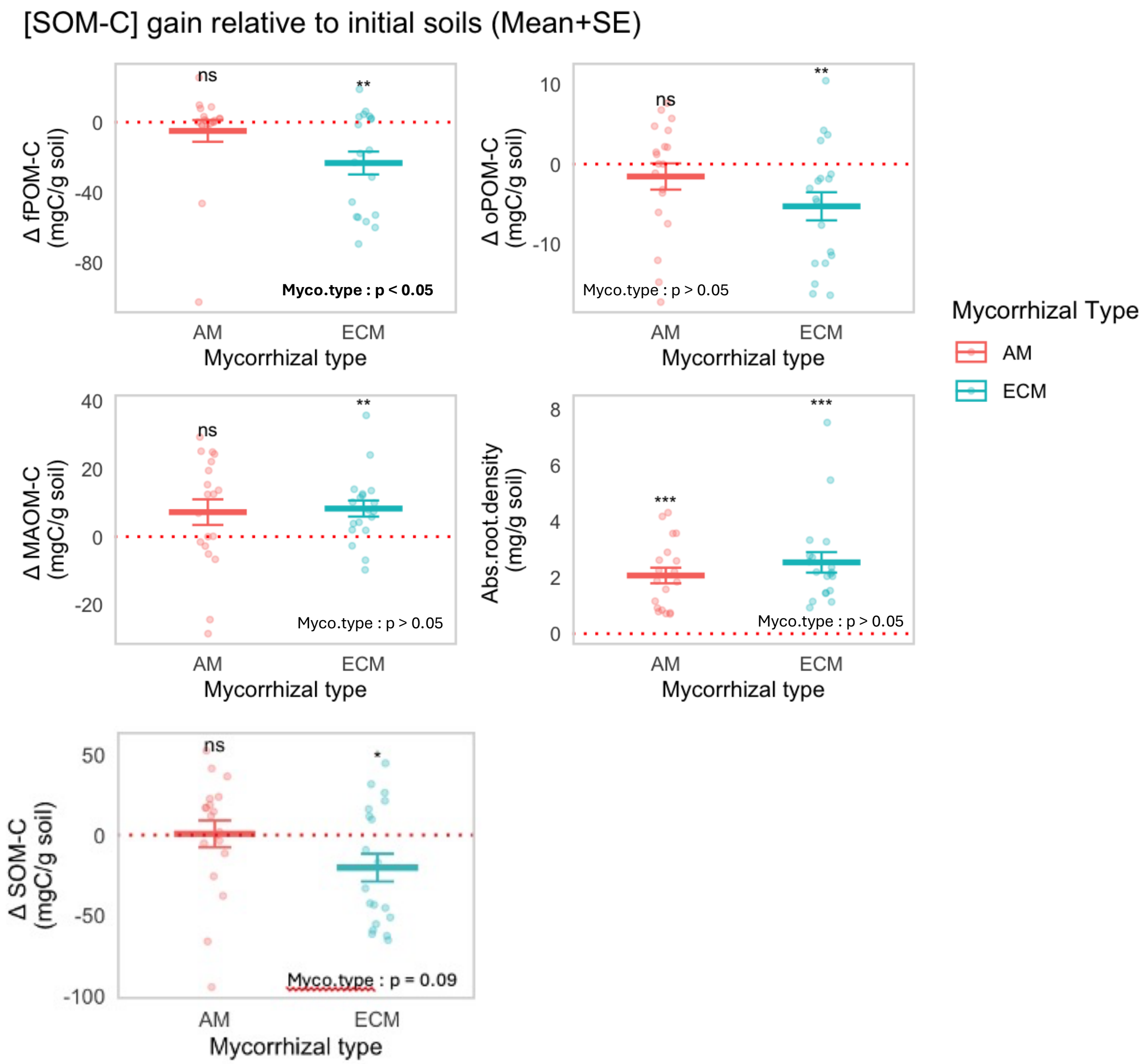


Fig S1) The effects of mycorrhizal type on changes in soil organic matter C concentrations (ΔmgC $g^{-1}$ soil) relative to initial soils across particulate and mineral-associated fractions, including free particulate organic matter (fPOM), occluded particulate organic matter (oPOM), and mineral-associated organic matter (MAOM), absorptive root density (mg.root $g^{-1}$ soil), and total soil organic matter (SOM) in 24 months of new root ingrowth. Points represent individual observations per each plot, and crossbars indicate the mean ± SE. Fixed effects of mycorrhizal type on the variables were evaluated using linear models (lm). Model summaries are shown within each panel.

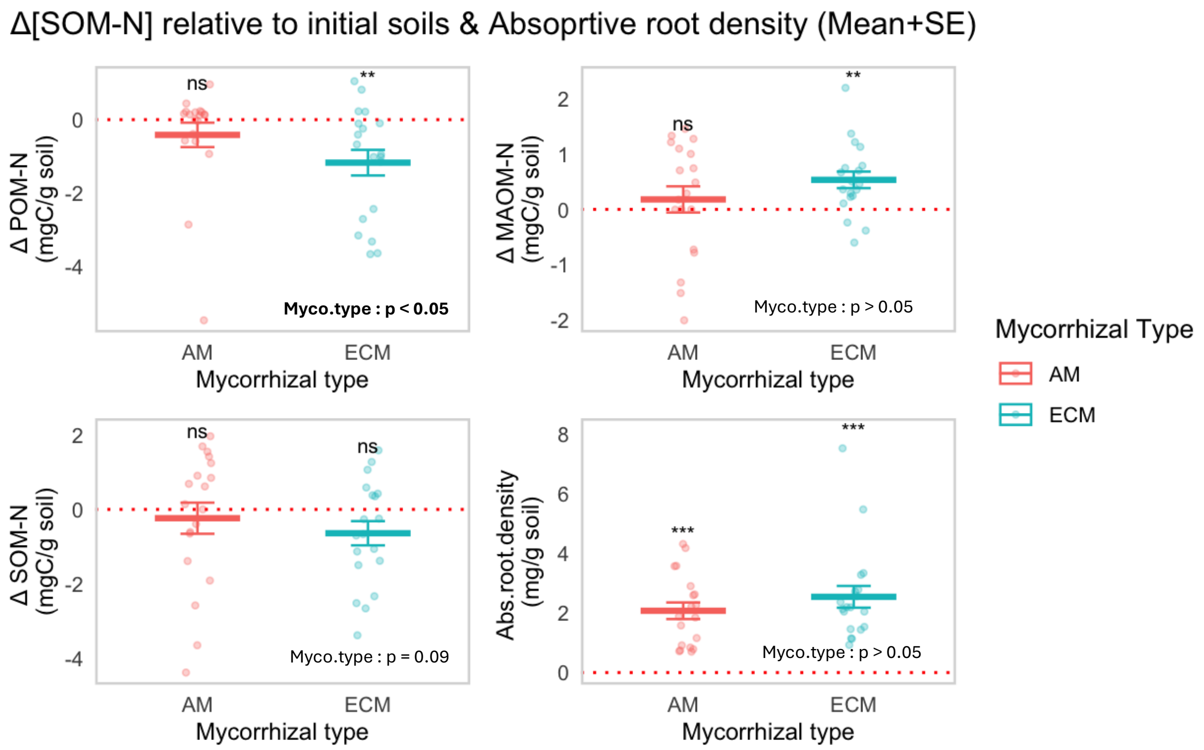


Fig S2) The effects of mycorrhizal type on changes in soil organic matter C concentrations (ΔmgC $g^{-1}$ soil) relative to initial soils across (a) particulate (POM) and (b) mineral-associated organic matter fractions (MAOM), (c) sum of all fraction (SOM) and (d) absorptive root density (mg.root $g^{-1}$ soil) in 24 months of new root ingrowth. Points represent individual observations per each plot, and crossbars indicate the mean ± SE. Fixed effects of mycorrhizal type on the variables were evaluated using linear models (lm). Model summaries are shown within each panel.

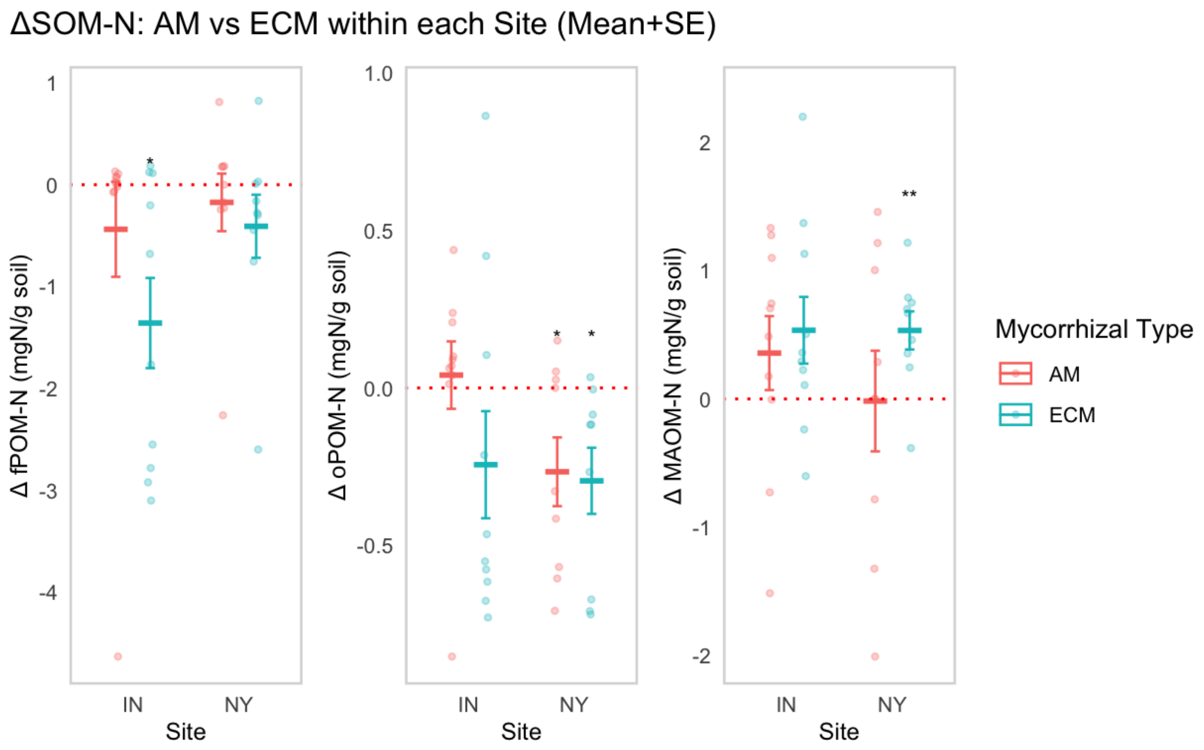


Fig. S3) The effects of Site, mycorrhizal type, and their interaction on changes in soil organic matter N concentrations relative to initial soils (ΔSOM-N; mg N $g^{-1}$ soil) in the free particulate organic matter (fPOM), occluded particulate organic matter (oPOM), and mineral-associated organic matter (MAOM) fractions in 24 months of new root ingrowth. Points represent individual observations, and crossbars indicate the mean ± SE for each Site × mycorrhizal type combination. Red dotted horizontal lines denote no change relative to initial soil (Δ = 0). Fixed effects of Site, mycorrhizal type, and their interaction were evaluated using linear models (lm). Asterisks indicate whether ΔSOM-C differed significantly from zero within each Site × mycorrhizal type combination ($P < 0.05$, $P < 0.01$), while model summaries are shown within each panel.

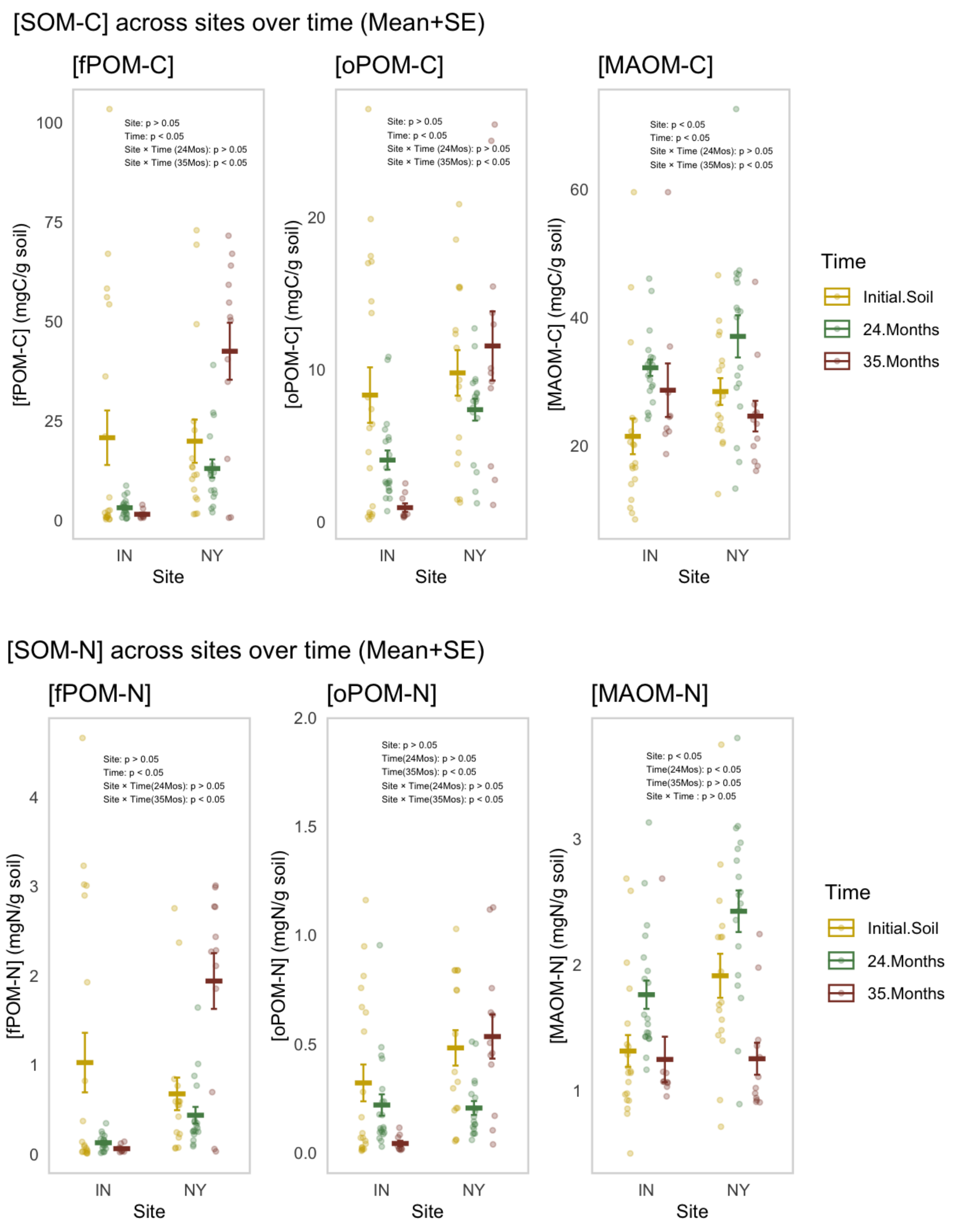


Fig. S4) The effects of Site, Time, and their interactions on soil organic matter C and N concentrations (mgN $g^{-1}$ soil) across particulate and mineral-associated fractions, including free particulate organic matter (fPOM), occluded particulate organic matter (oPOM), and mineral-associated organic matter (MAOM). Points represent individual observations, and crossbars indicate the mean ± SE for each Site × Time combination. Fixed effects of Site, Time, and their interaction were evaluated using linear mixed-effects models with Plot included as a random effect.

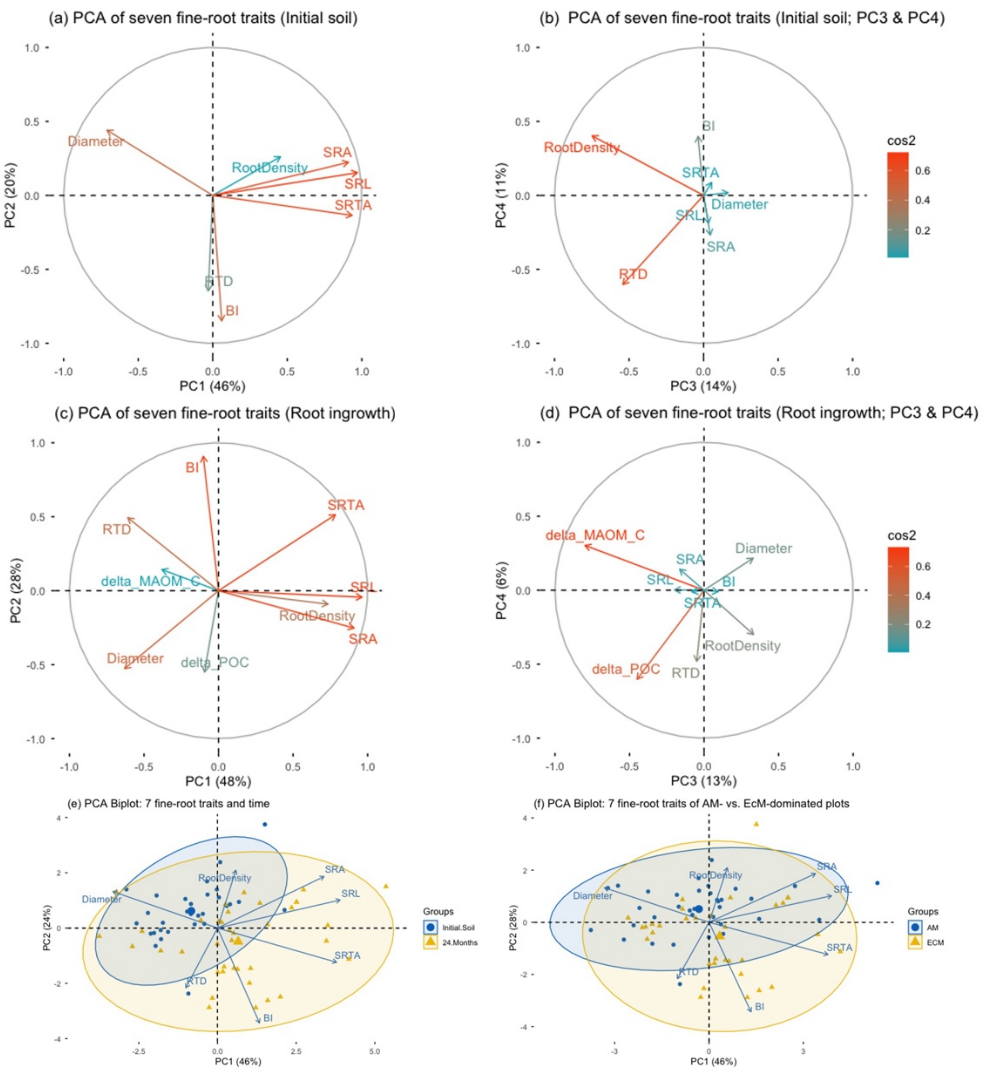


Fig. S5) Principal component analyses (PCA) of fine-root trait organization in initial soils and after two years of experimental root ingrowth across arbuscular mycorrhizal (AM)- and ectomycorrhizal (EcM)-dominated forests. (a, b) PCA correlation circles showing the relationships of seven fine-root traits with the first two principal components (PC1 and PC2; a) and the third and fourth components (PC3 and PC4; b) in initial soils (n = 30). (c, d) Corresponding PCA correlation circles for roots collected after two years of ingrowth (n = 38), including changes in particulate organic matter carbon (ΔPOM-C) and mineral-associated organic matter carbon (ΔMAOM-C) relative to initial soils. (e, f) PCA biplots showing the distributions of AM- and EcM-dominated plots in relation to root trait gradients at initial sampling (e) and after two years of ingrowth (f). Arrows indicate the direction and strength of variable correlations with the principal components, with colors in the correlation circles representing the quality of representation ($cos^2$). Points represent individual observations, with colors and shapes distinguishing mycorrhizal types; ellipses indicate 95% confidence regions. Separate PCAs were conducted for the two sampling stages to characterize stage-specific root trait organization. The proportion of variance explained by each component is indicated on the corresponding axes, and detailed PCA loadings are provided in Table S5.

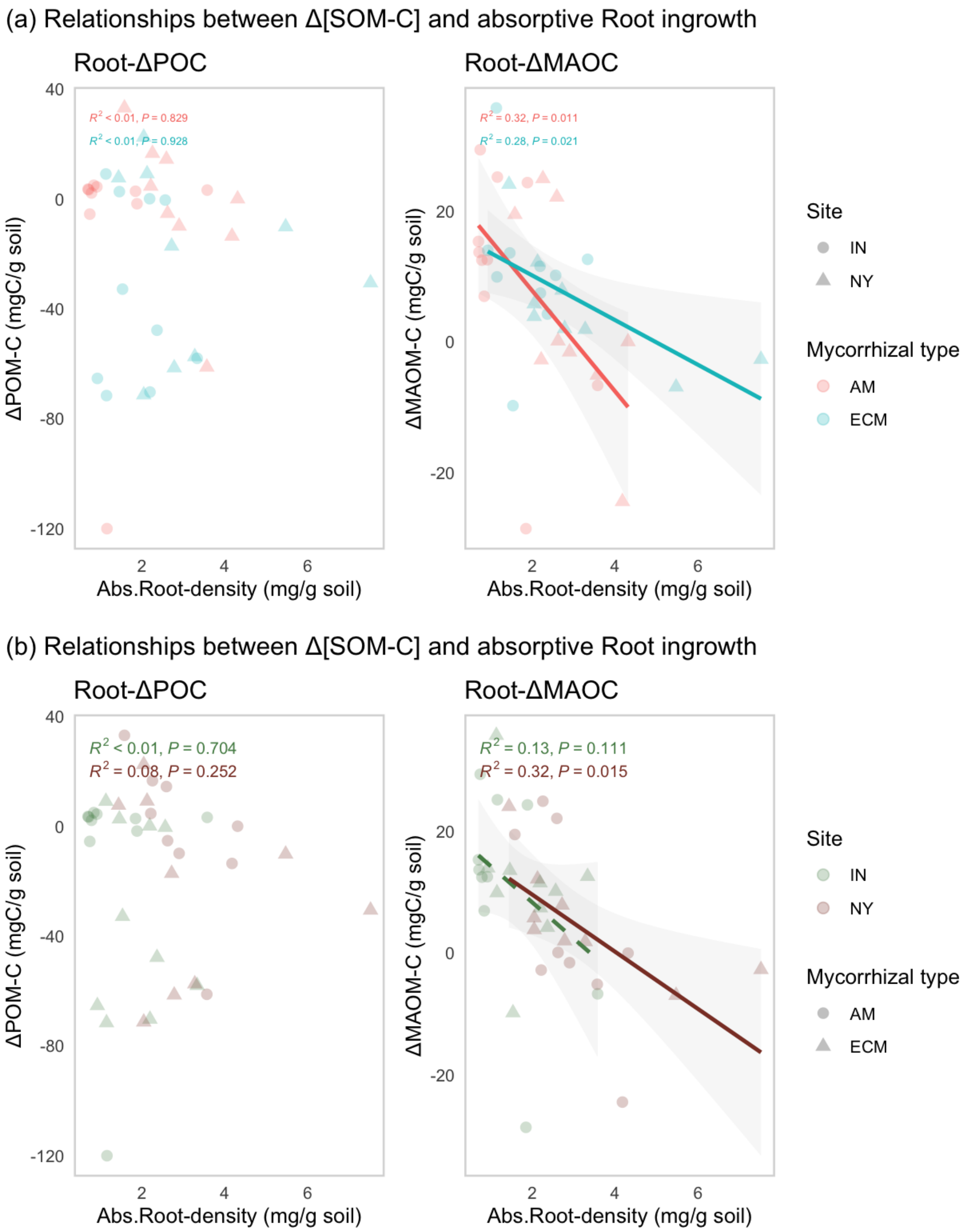


Fig. S6) Relationships between changes in soil organic matter C (ΔSOM-C) and absorptive root ingrowth density across sites and mycorrhizal types. Changes in particulate organic matter C (ΔPOM-C; sum of ΔC in the fPOM and oPOM fractions) and mineral-associated organic matter C (ΔMAOM-C) are expressed relative to initial soils. Points represent individual observations, with point shape indicating site (IN and NY) and color indicating mycorrhizal type (AM and ECM). Lines represent fitted linear regressions for each mycorrhizal type, with shaded areas indicating 95% confidence intervals. Model $R^2$ and P-values (a) for AM and ECM relationships and (b) for IN vs. NY are shown within each panel.

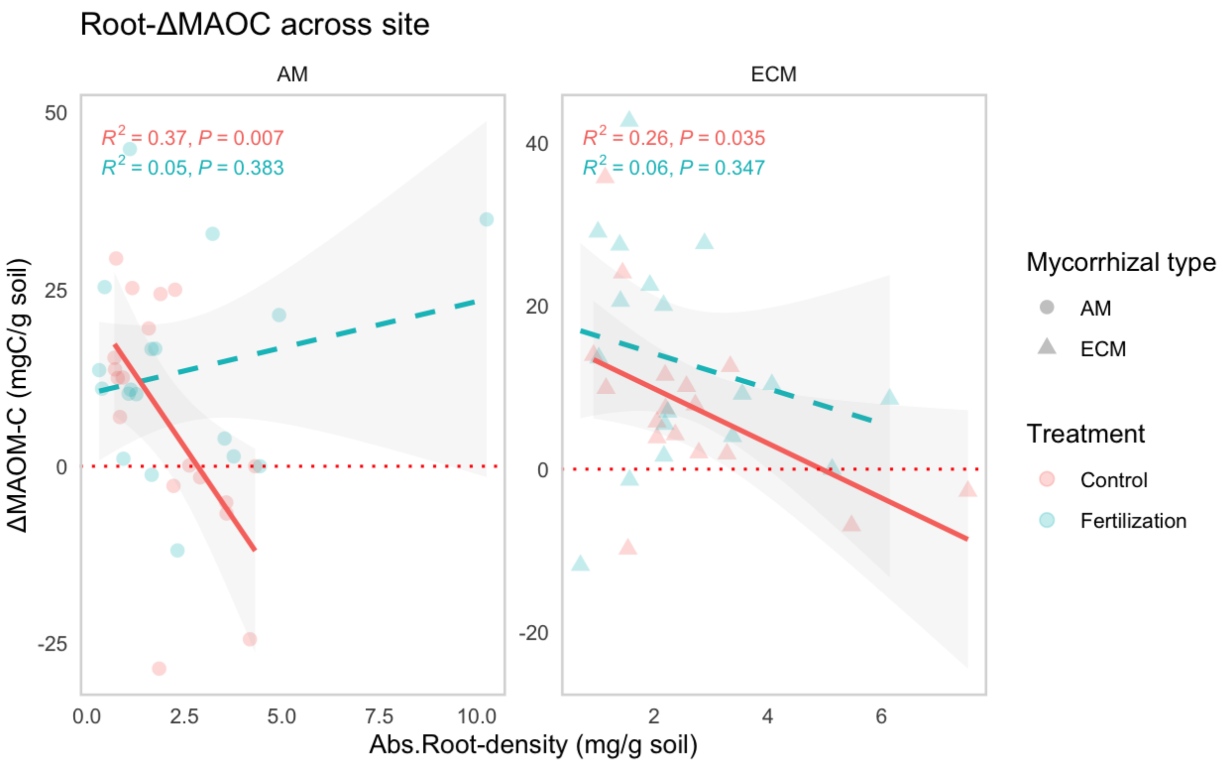


Fig. S7) Effects of fertilization on relationships between absorptive root ingrowth density and mineral-associated organic matter C formation (ΔMAOM-C) among AM vs. EcM tree associated plots. ΔMAOM-C represents the change in MAOM-C concentration relative to initial soils (mg C $g^{-1}$ soil). Points represent individual observations, with point shape indicating mycorrhizal type (AM and ECM) and color indicating treatment (control and fertilization). Lines represent treatment-specific linear regressions nested in different mycorrhizal type across sites, with shaded areas indicating 95% confidence intervals. Differences in root–MAOM relationships between control and fertilized cores were evaluated by testing the absorptive root density × treatment interaction. Model $R^2$ and P-values for treatment-specific relationships are shown within each panel.

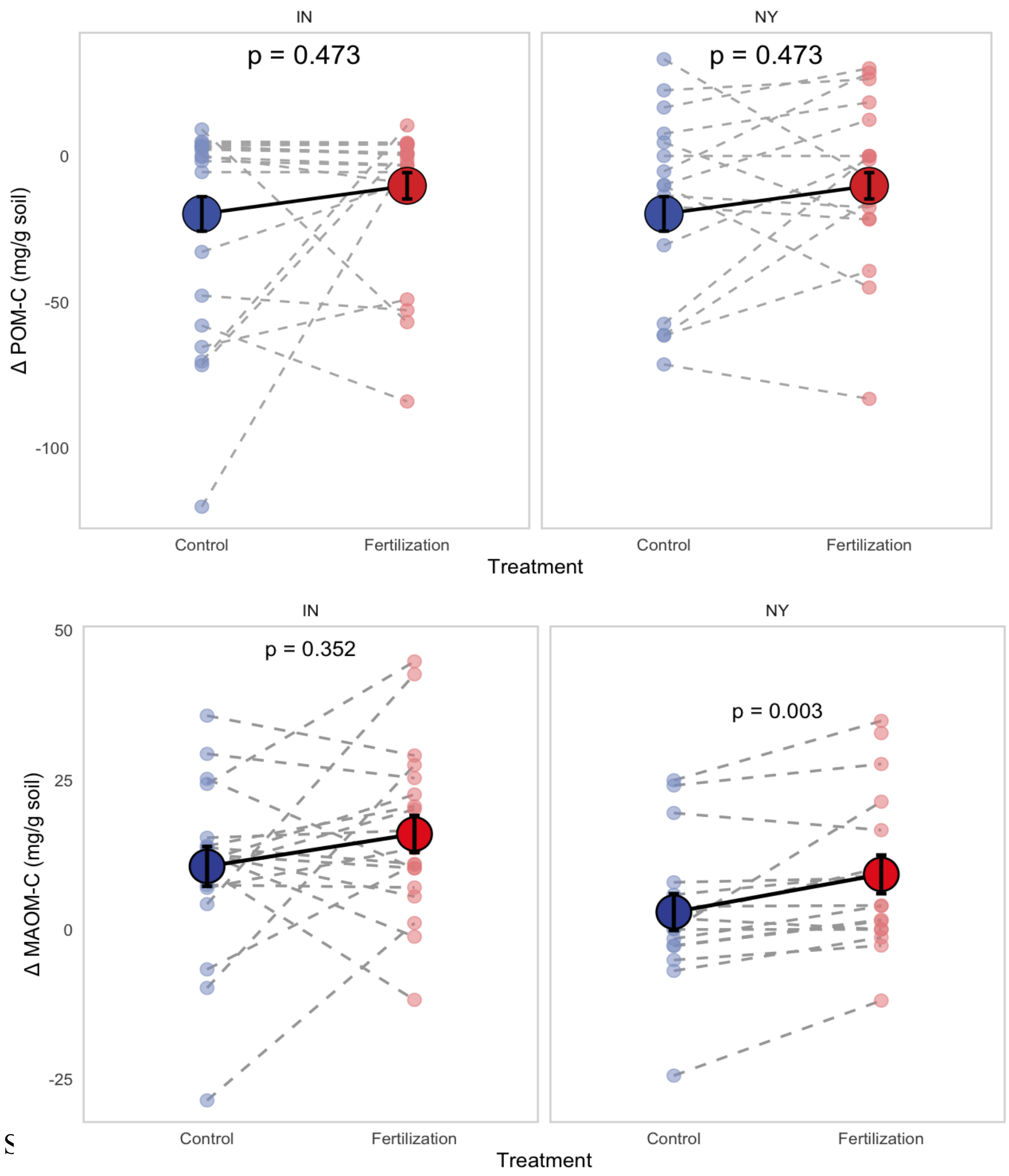


Fig. S[illegible] p panels) and mineral-associated organic matter C (ΔMAOM-C; bottom panels) relative to initial soils at IN and NY. Each point represents individual observations from control and fertilized cores, with dashed lines connecting paired observations from the same plot. Large points and solid lines represent treatment means ± SE. Treatment effects were evaluated using paired comparisons within each site, with P-values shown within each panel. Fertilization significantly increased ΔMAOM-C at NY ($P = 0.003$), whereas ΔMAOM-C at IN ($P = 0.352$) and ΔPOM-C at both IN and NY ($P = 0.473$) did not differ significantly between control and fertilized cores.

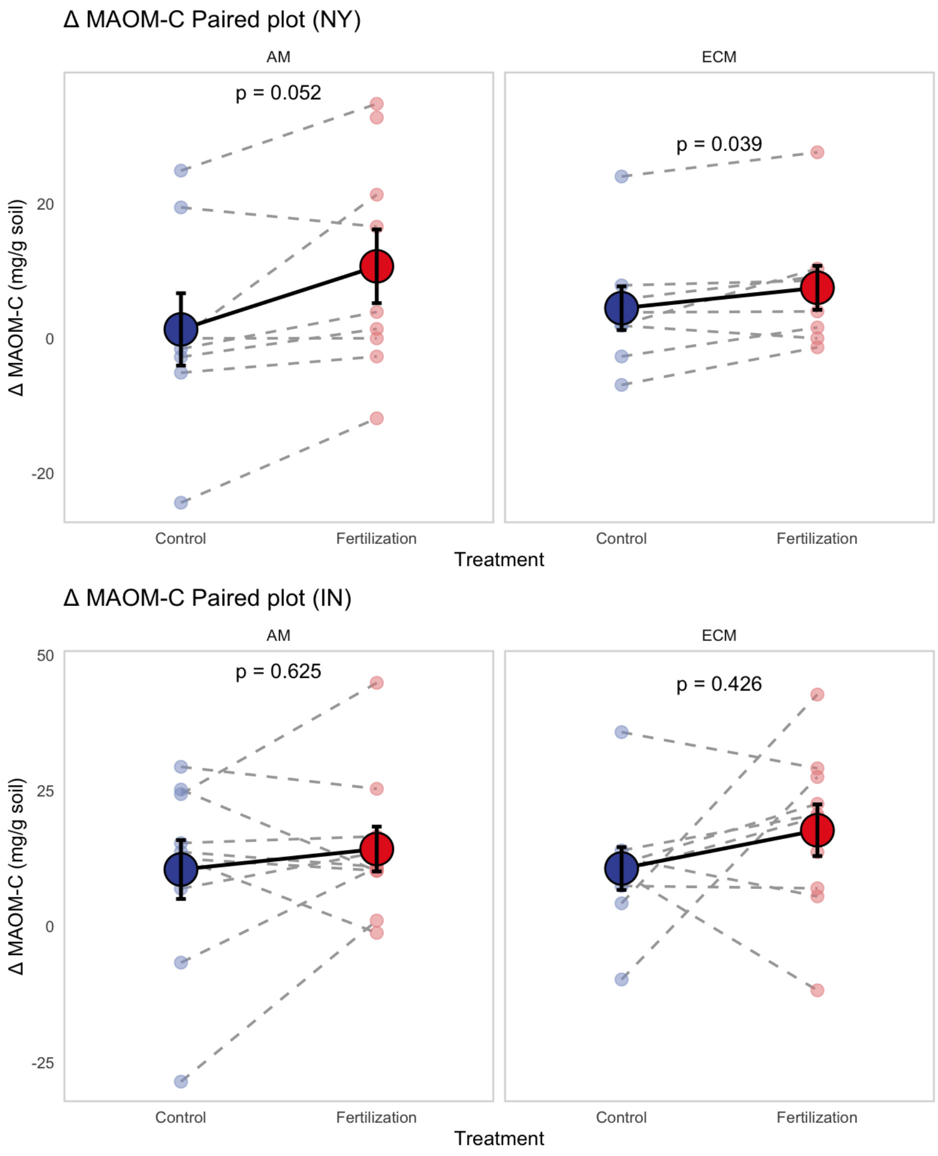


Fig. S9) Effects of fertilization on changes in mineral-associated organic matter C (ΔMAOM-C) relative to initial soils for AM- and ECM-associated plots at NY (top panels) and IN (bottom panels). Small points represent individual observations from control and fertilized cores, with dashed lines connecting paired observations from the same plot. Large points and error bars represent treatment means ± SE, and solid lines connect treatment means. Treatment effects were evaluated using paired comparisons separately for each mycorrhizal type within each site, with P-values shown within each panel. Fertilization significantly increased ΔMAOM-C in both AM ($p = 0.052$) and ECM ($p = 0.039$) plots at NY, whereas no significant fertilization effect was detected for AM ($p = 0.625$) or ECM ($p = 0.426$) plots at IN.

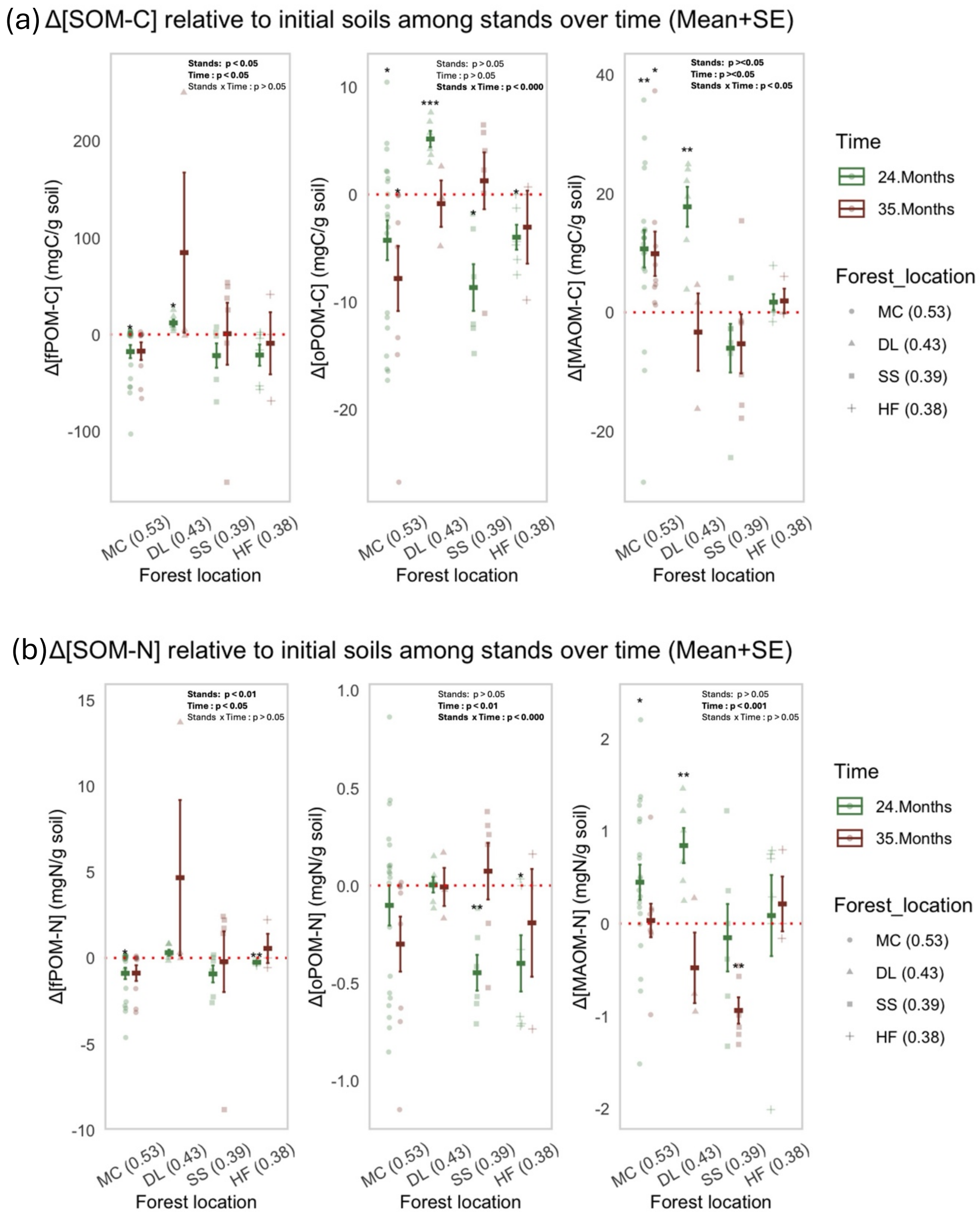


Fig. S10) The effects of forest location (stands), Time, and their interaction on changes in soil organic matter (a) C and (b) N concentrations relative to initial soils (ΔSOM-C; mg C $g^{-1}$ soil & ΔSOM-N; mg N $g^{-1}$ soil) in the free particulate organic matter (fPOM), occluded particulate organic matter (oPOM), and mineral-associated organic matter (MAOM) fractions. Each location represent MC: Moores Creek, IN, DL: Lake Goerge Wild Forest, NY, HF: Huntington Wildlife Forest, NY, and SS: Shingle Shanty Nature Preserve, NY. Points represent individual observations, and crossbars indicate the mean ± SE for each Site × Time combination. Red dotted horizontal lines denote no change relative to initial soil (Δ = 0). Fixed effects of

Site, Time, and their interaction were evaluated using linear mixed-effects models with Plot included as a random effect. Asterisks indicate whether ΔSOM-C differed significantly from zero within each Site × Time combination ($P < 0.05$, $P < 0.01$), while model summaries are shown within each panel.

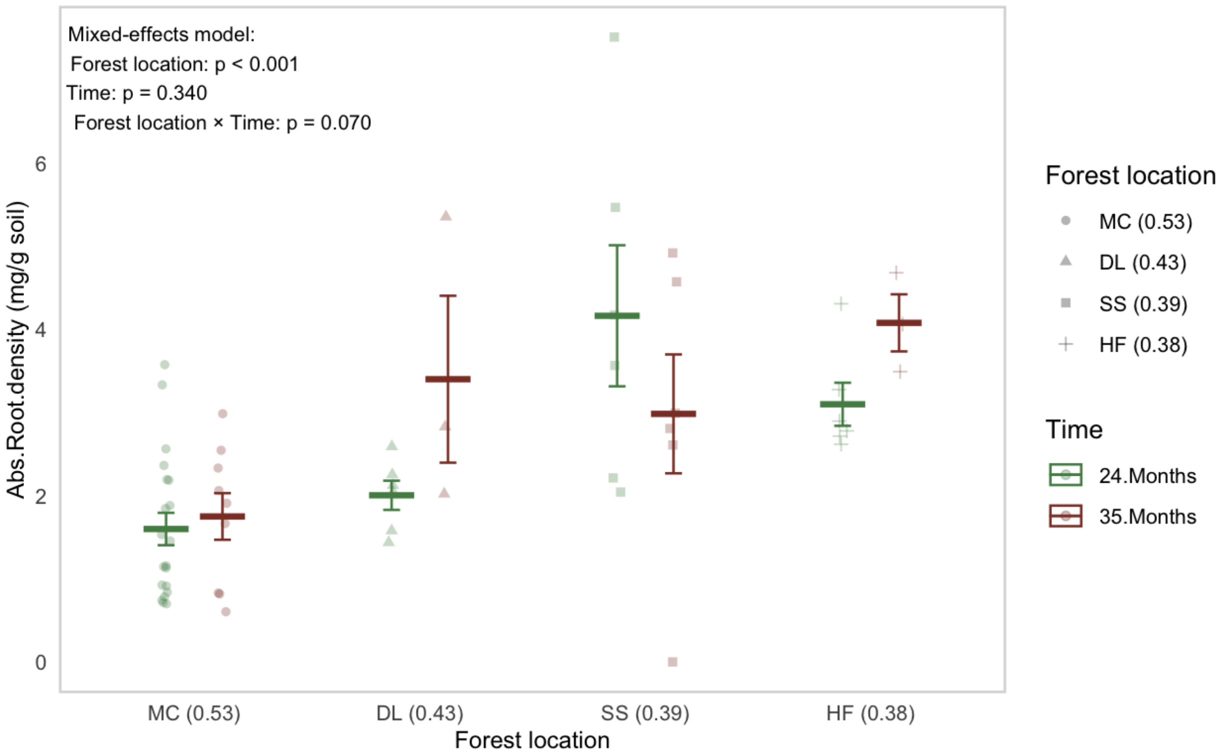


Fig. S15) The effects of forest location (stands), Time, and their interaction on absorptive root density ($g_{root}$ $g^{-1}$ soil ). Each location represents MC: Moores Creek, IN, DL: Lake Goerge Wild Forest, NY, HF: Huntington Wildlife Forest, NY, SS: Shingle Shanty Nature Preserve, NY. Points represent individual observations, and crossbars indicate the mean ± SE for each Site × Time combination. Red dotted horizontal lines denote no change relative to initial soil ($\Delta = 0$). Fixed effects of Site, Time, and their interaction were evaluated using linear mixed-effects models with Plot included as a random effect. Asterisks indicate whether ΔSOM-C differed significantly from zero within each Site × Time combination ($P < 0.05$, $P < 0.01$), while model summaries are shown within each panel.

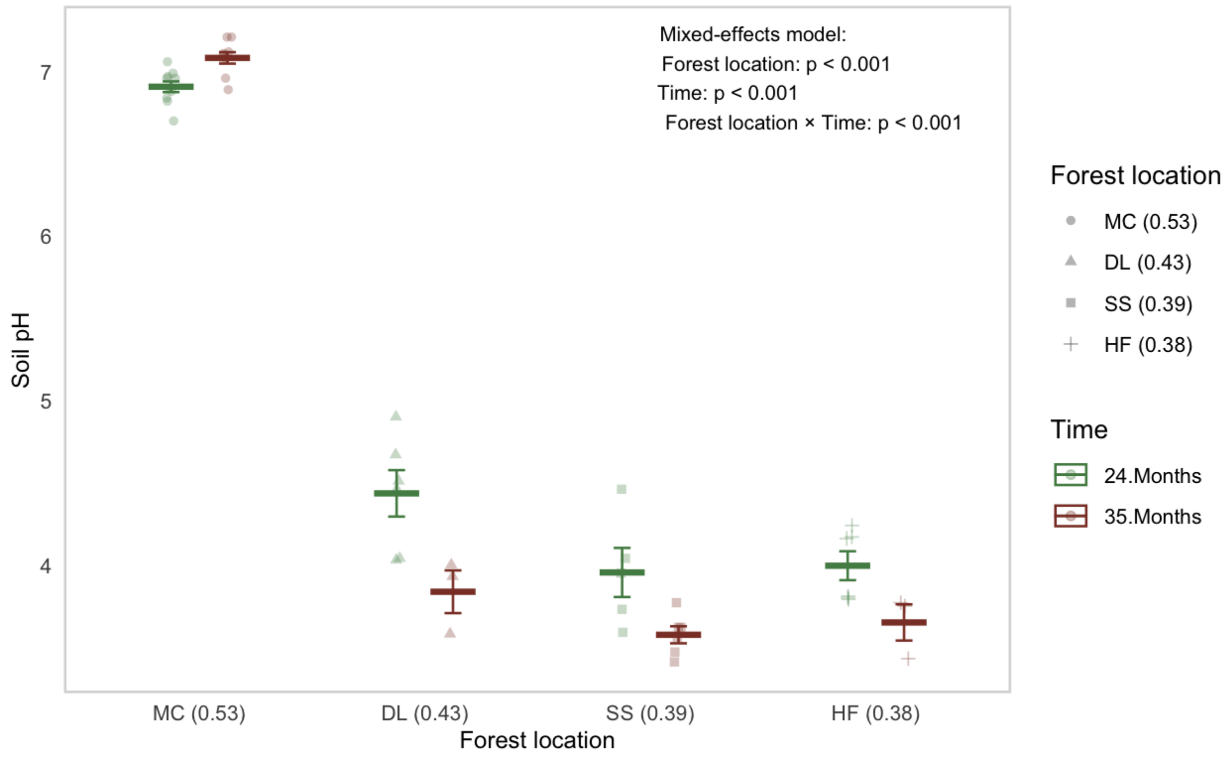


Fig. S16) The effects of forest location (stands), Time, and their interaction on soil pH. Each location represents MC: Moores Creek, IN, DL: Lake Goerge Wild Forest, NY, HF: Huntington Wildlife Forest, NY, SS: Shingle Shanty Nature Preserve, NY. Points represent individual observations, and crossbars indicate the mean ± SE for each Site × Time combination. Red dotted horizontal lines denote no change relative to initial soil ($\Delta = 0$). Fixed effects of Site, Time, and their interaction were evaluated using linear mixed-effects models with Plot included as a random effect. Asterisks indicate whether ΔSOM-C differed significantly from zero within each Site × Time combination ($P < 0.05$, $P < 0.01$), while model summaries are shown within each panel

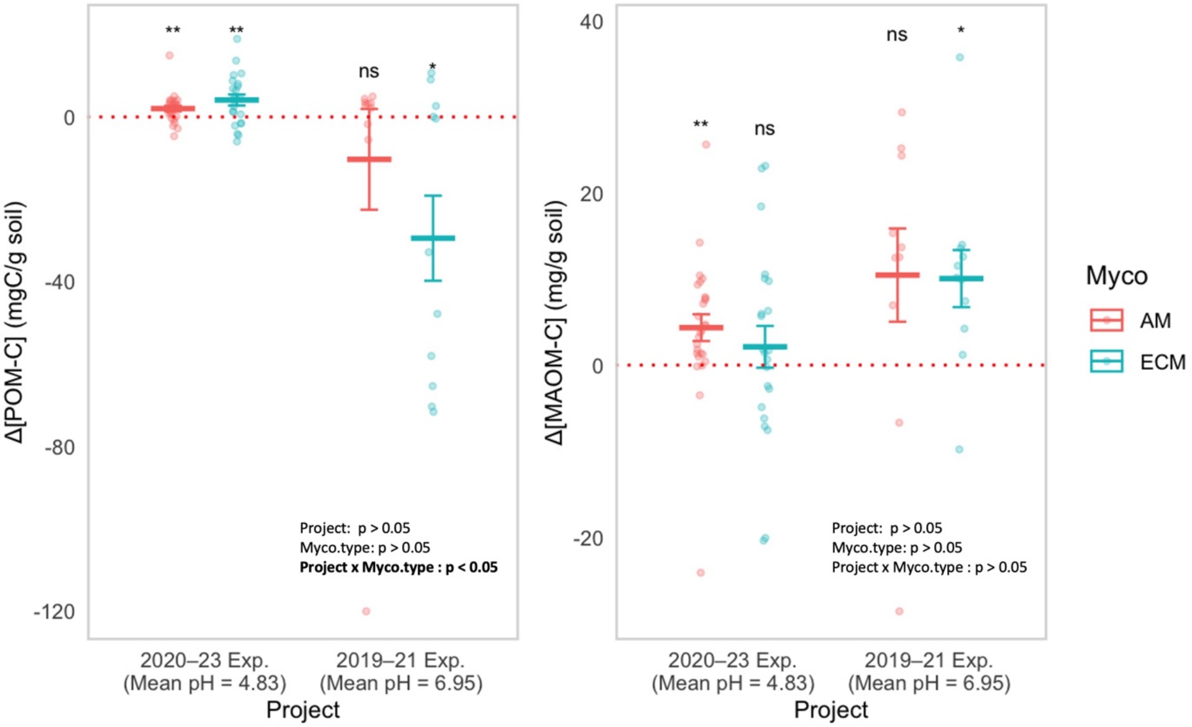


Fig. S17) Changes in particulate organic matter carbon (ΔPOM-C) and mineral-associated organic matter carbon (ΔMAOM-C) in AM- and EcM-dominated forests across two experiments with contrasting soil pH conditions. Changes in POM-C (left) and MAOM-C (right) were calculated relative to initial soil concentrations. The two experiments differed in average soil pH, with mean pH values of 4.83 (2020–2023 ingrowth experiment) and 6.95 (Our Experiment; 2019–2021). Points represent individual observations, with colors distinguishing arbuscular mycorrhizal (AM) and ectomycorrhizal (EcM) plots. Horizontal bars and error bars indicate means ± standard errors. Asterisks indicate significant differences from zero based on one-sample t-tests (*P < 0.05, **P < 0.01, ***P < 0.001), whereas ns indicates nonsignificant differences. Red dotted lines indicate zero change in SOM-C.

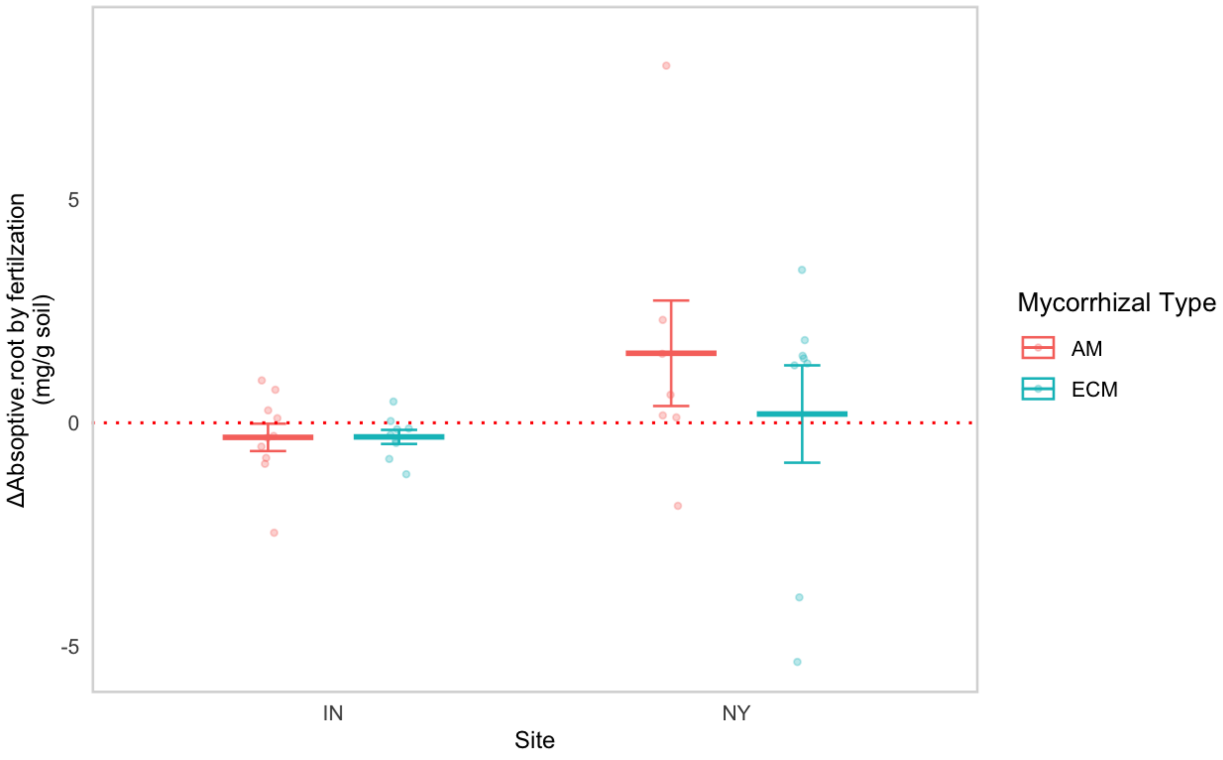


Fig. S18) Changes in absorptive root density (mg root $g^{-1}$ soil) due to fertilization in AM- and EcM-dominated forests across two experiments with contrasting soil pH conditions. Changes in absorptive root density were calculated relative to controls. Points represent individual observations, with colors distinguishing arbuscular mycorrhizal (AM) and ectomycorrhizal (EcM) plots. Horizontal bars and error bars indicate means ± standard errors. Changes in absorptive root density were found to be insignificant. Asterisks indicate significant differences from zero based on one-sample t-tests (*$P < 0.05$, **$P < 0.01$, ***$P < 0.001$), whereas ns indicates nonsignificant differences. Red dotted lines indicate zero change in SOM-C

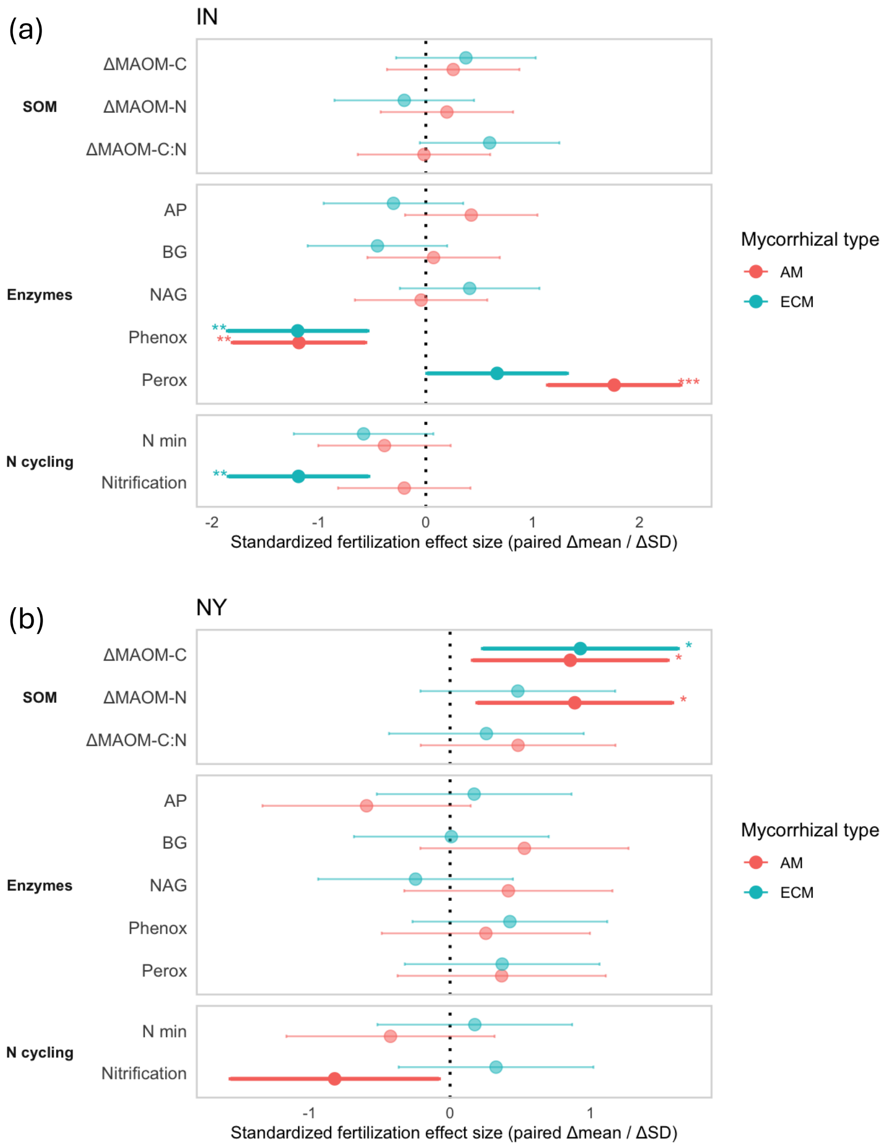


Fig. S19) Standardized effects of fertilization on soil organic matter dynamics, potential extracellular enzyme activities, and N cycling in AM- and EcM-dominated plots at (a) the Indiana (IN) site and (b) New York (NY) site. Fertilization effects are expressed as standardized paired effect sizes (Δmean/ΔSD) for changes in mineral-associated organic matter C (ΔMAOM-C), N (ΔMAOM-N), and C:N ratio (ΔMAOM-C:N); potential acid phosphatase (AP), β-glucosidase (BG), N-acetyl-β-glucosaminidase (NAG), phenol oxidase (Phenox), and peroxidase (Perox) activities; and net N mineralization (N min) and nitrification rates. Points represent mean standardized fertilization effects and Points represent standardized mean paired differences and horizontal bars indicate approximate 95% confidence intervals. Positive values indicate greater values under fertilization relative to paired controls, whereas negative values indicate lower values under fertilization. The vertical dotted line denotes no fertilization effect (effect size = 0). Colors distinguish AM- and EcM-dominated plots. Asterisks denote significant paired fertilization effects ($P < 0.05$, $^{*}P < 0.01$, $^{**}P < 0.001$).

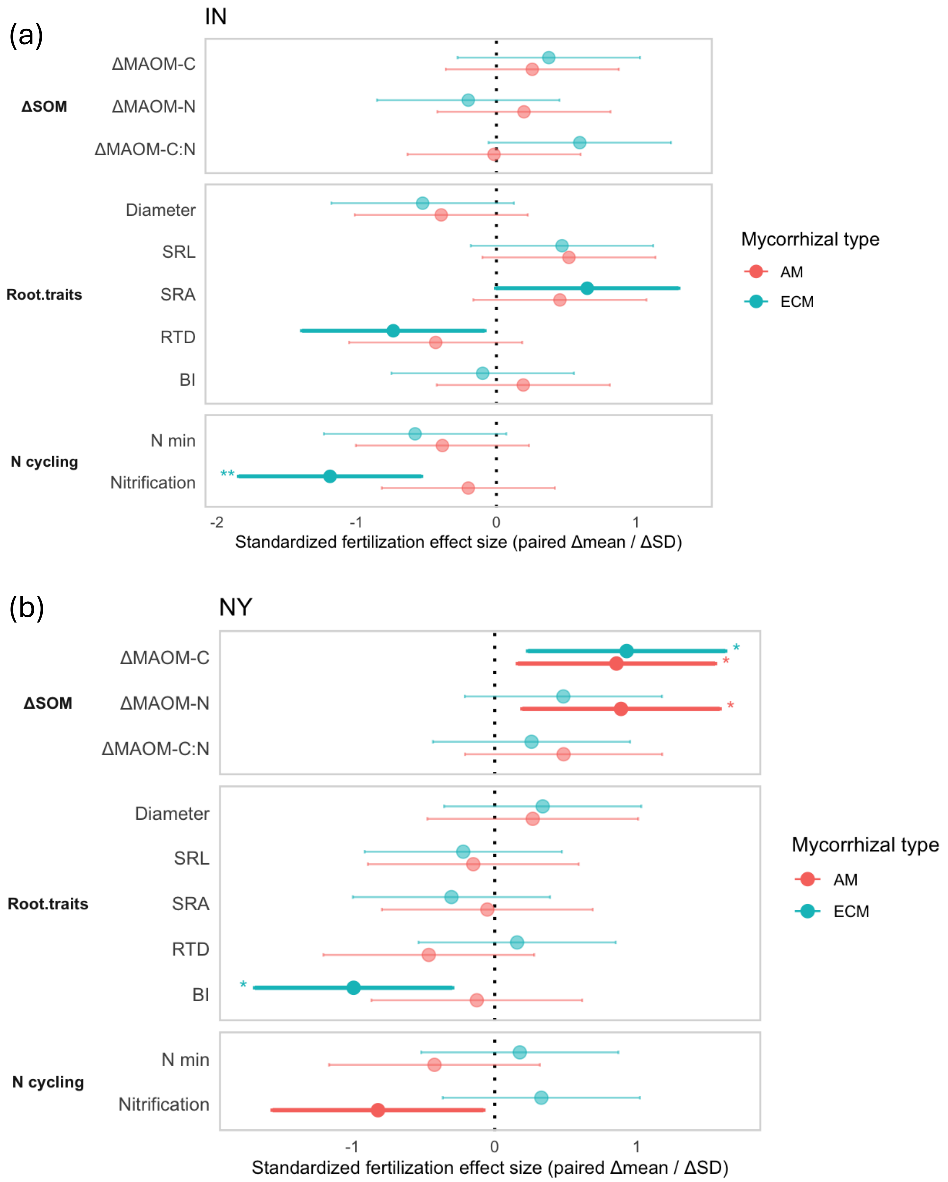


Fig. S20) Standardized effects of fertilization on mineral-associated organic matter (MAOM) dynamics, root functional traits, and N cycling in AM- and EcM-dominated plots at (a) the Indiana (IN) site and (b) New York (NY) site.. Fertilization effects are expressed as standardized paired effect sizes (paired mean difference/SD of paired differences) for changes in MAOM-C (ΔMAOM-C), MAOM-N (ΔMAOM-N), and MAOM C:N (ΔMAOM-C:N); root diameter, specific root length (SRL), specific root area (SRA), root tissue density (RTD), and branching intensity (BI); and net N mineralization (N min) and nitrification rates. Points represent standardized paired mean differences, and horizontal bars indicate approximate 95% confidence intervals. Positive values indicate greater values under fertilization relative to paired controls, whereas negative values indicate lower values under fertilization. The vertical dotted line indicates no fertilization effect. Colors distinguish AM- and EcM-dominated plots. Asterisks indicate significant paired fertilization effects ($P < 0.05$, $*P < 0.01$, $**P < 0.001$).

**Table S1 ) Linear models showed that the effect of dominant tree mycorrhizal type and sites on the changes in the concentrations of soil organic matter C and N (mg of C per g of soil, Free particulate organic matter: fPOM C, Occluded particulate organic matter: oPOM C, mineral-associated and particulate organic matter: MAOM C) and absorptive root density (mg of absorptive root per g of soil).**

| Predictors | ΔFree particulate organic matter C concentration (mg C of fPOM / g soil) | | | ΔOccluded particulate organic matter C concentration (mg C of oPOM / g soil) | | | ΔMineral-associated C concentration (mg C of MAOM / g soil) | | | Absorptive root density (mg of roots / g soil) | | |
|---|---|---|---|---|---|---|---|---|---|---|---|---|
| | Estimate | SE | p | Estimate | SE | p | Estimate | SE | p | Estimate | SE | p |
| Intercept | 0.851 | 2.561 | 0.742 | 2.189 | 2.740 | 0.430 | 30.999 | 2.495 | **0.000** | 1.315 | 0.388 | **0.002** |
| Mycorrhizal type (ECM) | -0.110 | 3.622 | 0.976 | -0.115 | 3.875 | 0.977 | -8.662 | 3.528 | **0.020** | 0.568 | 0.549 | 0.309 |
| Site (NY) | 4.864 | 3.721 | 0.200 | 1.148 | 3.981 | 0.775 | -20.952 | 3.625 | **0.000** | 1.323 | 0.605 | **0.036** |
| Mycorrhizal type × Site | -8.211 | 5.348 | 0.134 | -1.589 | 5.722 | 0.783 | 7.371 | 5.210 | 0.167 | 0.137 | 0.828 | 0.869 |
| Adj R-squared | Adj R-squared = 0.0429 | | | Adj R-squared = -0.08484 | | | **Adj R-squared = 0.5661*** | | | **Adj R-squared = 0.2475**** | | |

| Predictors | ΔFree particulate organic matter N concentration (mg N of fPOM / g soil) | | | ΔOccluded particulate organic matter N concentration (mg N of oPOM / g soil) | | | ΔMineral-associated N concentration (mg N of MAOM / g soil) | | |
|---|---|---|---|---|---|---|---|---|---|
| | Estimate | SE | p | Estimate | SE | p | Estimate | SE | p |
| Intercept | -0.870 | 2.056 | 0.675 | 0.172 | 0.145 | 0.243 | 1.463 | 0.310 | **0.000** |
| Mycorrhizal type (ECM) | -2.743 | 2.907 | 0.352 | 0.017 | 0.205 | 0.933 | -0.308 | 0.438 | 0.487 |
| Site (NY) | -2.121 | 2.987 | 0.483 | -0.128 | 0.210 | 0.548 | -0.433 | 0.450 | 0.343 |
| Mycorrhizal type × Site | -9.120 | 4.293 | **0.041** | -0.116 | 0.302 | 0.705 | -0.121 | 0.646 | 0.853 |
| Adj R-squared | **Adj R-squared = 0.3653**** | | | Adj R-squared = 0.03852 | | | Adj R-squared = 0.01431 | | |

**Table S2 ) Linear mixed-effect models describing the effect of absorptive root ingrowth and dominant tree mycorrhizal type on the changes in the concentrations of mineral-associated organic matter C and N. Study sites (IN vs. NY) were set as a random effect to account for differences in CDI, soil order, pH, and other site-level variation.**

| Predictors | Δ MAOM-C (mg C of MAOM / g soil) | | | Δ MAOM-N (mg N of MAOM / g soil) | | |
|---|---|---|---|---|---|---|
| | Estimate | SE | p | Estimate | SE | p |
| Intercept | 30.761 | 7.314 | **0.007** | 1.744 | 0.459 | **0.002** |
| Abs. root density | -5.810 | 2.339 | **0.018** | -0.321 | 0.208 | 0.133 |
| Mycorrhizal type (ECM) | -14.137 | 6.132 | **0.027** | -0.860 | 0.579 | 0.147 |
| Abs. root density × Mycorrhizal type | 4.772 | 2.556 | 0.070 | 0.282 | 0.240 | 0.248 |
| % explained by Random-effects | | | **40.1%**** | | | 3.20% |

**Table S3) Linear mixed-effects models summarizing the effects of Site, Time, and their interaction on soil organic matter C and N concentrations (mg $g^{-1}$ soil) across particulate and mineral-associated fractions, including free particulate organic matter (fPOM), occluded particulate organic matter (oPOM), and mineral-associated organic matter (MAOM). Models were fitted with Site, Time, and their interaction as fixed effects and Plot as a random effect.**

| Predictors | Free particulate organic matter C concentration (mg C of fPOM / g soil) | | | Occluded particulate organic matter C concentration (mg C of oPOM / g soil) | | | Mineral-associated C concentration (mg C of MAOM / g soil) | | |
|---|---|---|---|---|---|---|---|---|---|
| | Estimate | SE | p | Estimate | SE | p | Estimate | SE | p |
| Intercept | 20.698 | 4.234 | **0.000** | 8.331 | 1.222 | **0.000** | 21.444 | 2.289 | **0.000** |
| Site (NY) | -0.871 | 6.247 | 0.889 | 1.520 | 1.802 | 0.401 | 6.998 | 3.374 | **0.041** |
| Time (24 months) | -17.615 | 5.988 | **0.004** | -4.266 | 1.664 | **0.013** | 10.691 | 2.933 | **0.001** |
| Time (35 months) | -19.257 | 7.601 | **0.013** | -7.460 | 2.139 | **0.001** | 7.788 | 3.829 | **0.046** |
| Site × Time (NY × 24 months) | 10.740 | 8.768 | 0.224 | 1.788 | 2.437 | 0.466 | -2.111 | 4.300 | 0.625 |
| Site × Time (NY × 35 months) | 41.884 | 10.428 | **0.000** | 9.194 | 2.928 | **0.002** | -10.827 | 5.229 | **0.042** |
| % explained by Random-effects (plot) | | | 0.0% | | | 7.3% | | | 17.9% |

| Predictors | Free particulate organic matter N concentration (mg N of fPOM / g soil) | | | Occluded particulate organic matter N concentration (mg N of oPOM / g soil) | | | Mineral-associated N concentration (mg N of MAOM / g soil) | | |
|---|---|---|---|---|---|---|---|---|---|
| | Estimate | SE | p | Estimate | SE | p | Estimate | SE | p |
| Intercept | 1.023 | 0.187 | **0.000** | 0.319 | 0.061 | **0.000** | 1.315 | 0.129 | **0.000** |
| Site (NY) | -0.347 | 0.276 | 0.211 | 0.164 | 0.090 | 0.072 | 0.596 | 0.190 | **0.002** |
| Time (24 months) | -0.898 | 0.256 | **0.001** | -0.102 | 0.082 | 0.222 | 0.447 | 0.179 | **0.016** |
| Time (35 months) | -0.960 | 0.329 | **0.005** | -0.278 | 0.106 | **0.010** | -0.063 | 0.229 | 0.783 |
| Site × Time (NY × 24 months) | 0.655 | 0.376 | 0.086 | -0.177 | 0.120 | 0.146 | 0.065 | 0.263 | 0.805 |
| Site × Time (NY × 35 months) | 2.211 | 0.450 | **0.000** | 0.330 | 0.145 | **0.026** | -0.581 | 0.314 | 0.068 |
| % explained by Random-effects (plot) | | | 5.8% | | | 8.9% | | | 3.4% |

**Table S4 ) Linear models showed that the effect of dominant tree mycorrhizal type and sites on the changes in the concentrations of soil organic matter C and N (Δmg of C per g of soil, Particulate organic matter: total POM C (fPOM-C + oPOM-C), mineral-associated and particulate organic matter: MAOM C) and two-year absorptive root ingrowth (mg of absorptive root per g of soil).**

| Predictors | Absorptive root density (mg of roots / g soil) | | | Δtotal.Particulate organic matter C concentration (mg C of POM / g soil) | | | ΔMineral-associated C concentration (mg C of MAOM / g soil) | | |
|---|---|---|---|---|---|---|---|---|---|
| | Estimate | SE | p | Estimate | SE | p | Estimate | SE | p |
| Intercept | 1.315 | 0.391 | **0.002** | 3.040 | 4.100 | 0.464 | 30.999 | 2.495 | **0.000** |
| Mycorrhizal type (ECM) | 1.515 | 0.586 | **0.014** | 6.013 | 5.957 | 0.320 | -20.952 | 3.625 | **0.000** |
| Site (NY) | 0.568 | 0.552 | 0.311 | -0.225 | 5.799 | 0.969 | -8.662 | 3.528 | **0.020** |
| Mycorrhizal type × Site | -0.055 | 0.816 | 0.947 | -9.800 | 8.563 | 0.261 | 7.371 | 5.210 | 0.167 |
| Adj R-squared | **Adj R-squared = 0.256**** | | | Adj R-squared = -0.01005 | | | **Adj R-squared = 0.5661***** | | |

**Table S5)** Results of principal component analysis (PCA) on the seven root traits in the initial soils and 2 year root-ingrowth core soils measured at Moores Creek, IN, and Adirondacks regions, NY, USA. The table includes the proportion of variation explained (top section) and loading scores of traits on each component (bottom section).

| Analyses | | PC 1 | PC2 | PC3 | PC4 |
|---|---|---|---|---|---|
| | Eigenvalue | 3.36 | 1.49 | 0.91 | 0.76 |
| | Variance | 0.48 | 0.21 | 0.13 | 0.11 |
| PCA of 7 root traits of initial soils (n=30; Fig. S5a & S5b) | Root Density | 0.25 | 0.21 | **0.65** | -0.67 |
| | Root Diameter | -0.39 | 0.36 | -0.21 | -0.08 |
| | SRL | **0.53** | 0.13 | -0.16 | 0.03 |
| | SRA | **0.50** | 0.18 | -0.26 | -0.04 |
| | RTD | -0.02 | **-0.53** | **-0.52** | **-0.66** |
| | BI | 0.03 | **-0.69** | 0.40 | 0.22 |
| | SRTA | **0.51** | -0.11 | -0.10 | 0.23 |
| | Eigenvalue | 3.84 | 2.00 | 1.14 | 0.84 |
| | Variance | 0.43 | 22.30 | 12.70 | 9.36 |
| PCA of 7 root traits & ΔSOM-C in 2 years (n=38; ; Fig. S5c & S5d) | Root Density | -0.37 | -0.06 | 0.31 | -0.33 |
| | Root Diameter | -0.32 | -0.37 | 0.31 | 0.24 |
| | SRL | **0.49** | -0.03 | -0.19 | 0.00 |
| | SRA | **0.46** | -0.18 | -0.15 | 0.15 |
| | RTD | -0.31 | 0.35 | -0.05 | **-0.52** |
| | BI | -0.31 | 0.35 | -0.05 | **-0.52** |
| | SRTA | **0.40** | 0.36 | -0.08 | -0.01 |
| | ΔPOM-C | -0.05 | -0.39 | -0.42 | **-0.66** |
| | ΔMAOM-C | -0.19 | 0.10 | **-0.75** | 0.33 |
| | Eigenvalue | 4.06 | 1.33 | 0.86 | 0.47 |
| | Variance | 0.58 | 0.19 | 0.12 | 0.67 |
| PCA of 7 root traits of 2 years ingrowth core soils in AM plots acorss site (n=19; Fig. 3a) | Root Density | 0.30 | **0.49** | -0.09 | **-0.79** |
| | Root Diameter | -0.37 | 0.01 | **0.57** | -0.35 |
| | SRL | **0.49** | -0.01 | -0.06 | 0.10 |
| | SRA | **0.48** | -0.03 | 0.06 | 0.14 |
| | RTD | -0.30 | -0.02 | -0.81 | -0.14 |
| | BI | 0.00 | **-0.82** | -0.03 | **-0.44** |
| | SRTA | **0.46** | -0.29 | -0.01 | -0.11 |
| | Eigenvalue | 4.22 | 1.55 | 0.71 | 0.37 |
| | Variance | 0.60 | 0.22 | 0.10 | 0.52 |
| PCA of 7 root traits of 2 years ingrowth core soils in EcM plots acorss site (n=19; Fig. 3a) | Root Density | **0.40** | -0.06 | -0.30 | **-0.83** |
| | Root Diameter | -0.27 | **-0.47** | **-0.68** | 0.05 |
| | SRL | **0.48** | 0.12 | 0.06 | 0.08 |
| | SRA | **0.47** | -0.11 | -0.12 | 0.13 |
| | RTD | **-0.40** | 0.27 | 0.30 | **-0.50** |
| | BI | 0.32 | **0.50** | -0.50 | 0.00 |
| | SRTA | 0.24 | **0.65** | -0.30 | 0.17 |

Note) PCs with eigenvalue close to or greater than 1 are considered as significant (Tabachnik & Fidell, 1996). Variable loading scores greater than 0.4 on each component appear in bold. SRA, specific root area ($cm^2\ g^{-1}$); specific root length ($m\ g^{-1}$); Diameter; fine-root mean diameter (cm); RTD, root tissue density ($mg\ cm^{-3}$); BI, branching intensity (tips per cm), specific root tip abundance (SRTA): # of root tips x $mass^{-1}$, ΔPOM-C (sum of ΔfPOM and ΔoPOM; mgC of SOM C & N $g^{-1}$ soil), and ΔMAOM-C (mgC of SOM C $g^{-1}$ soil).

**Table S6) Means and percent changes of soil organic matter C and N concentrations (mg $g^{-1}$ soil) across particulate and mineral-associated fractions of initial vs. 2 year root ingrowth core soils, including free particulate organic matter (fPOM), occluded particulate organic matter (oPOM), and mineral-associated organic matter (MAOM)**

| Site / mycorrhizal types | | Initial fPOM-C (mg C of fPOM / g soil) | 24 Month fPOM-C (mg C of fPOM / g soil) | Percen fPOM_C (%) | Initial oPOM-C (mg C of oPOM / g soil) | 24 Month oPOM-C (mg C of oPOM / g soil) | Percen oPOM_C (%) | Initial MAOM-C (mg C of MAOM / g soil) | 24 Month MAOM-C (mg C of MAOM / g soil) | Percen MAOM-C (%) |
|---|---|---|---|---|---|---|---|---|---|---|
| | n | Means | Means | Means | | | | Means | Means | Means |
| Time (Initial soil) | | | | | | | | | | |
| Moores Creek (IN) | 20 | | | | | | | | | |
| AM plots | 10 | 11.14 | 1.44 | -87.03 | 3.92 | 3.32 | -15.4 | 22.39 | 32.84 | 46.68 |
| EcM plots | 10 | 27.51 | 4.72 | -82.84 | 11.58 | 4.82 | -58.44 | 18.63 | 31.43 | 68.66 |
| Adirondack (ADK) (NY) | 18 | | | | | | | | | |
| AM plots | 9 | 12.76 | 12.24 | -4.03 | 10.05 | 7.61 | -24.32 | 29.93 | 35.35 | 18.11 |
| EcM plots | 9 | 27.31 | 13.66 | -49.98 | 9.1 | 7.14 | -21.54 | 32.65 | 38.69 | 18.51 |
